# Focal drug uncaging for the precise control of endogenous nicotinic receptors in behaving mice

**DOI:** 10.64898/2026.10.02.755724

**Authors:** Nicolas Guyon, Monir Mohayyaei, Antoine Bergel, Gawain Grellier, Joachim Jehl, Nicolas Pietrancosta, Nelson Rebola, Amaury François, Philippe Faure, Graham CR Ellis-Davies, Alexandre Mourot

**Affiliations:** Laboratoire Plasticité du Cerveau, ESPCI Paris, Université Paris Sciences et Lettres, CNRS, Paris, France; Department of Neuroscience, Karolinska Institutet, Stockholm, Sweden; Institut du Cerveau - Paris Brain Institute, Sorbonne Université, CNRS, Inserm, Paris, France; Institut de Génomique Fonctionnelle (IGF), Université de Montpellier, CNRS, Inserm, Montpellier, France; Laboratoire des Biomolécules, École Normale Supérieure, Université Paris Sciences et Lettres, CNRS, Paris, France; Neuroscience Paris Seine - Institut de Biologie Paris Seine (NPS - IBPS), Sorbonne Université, Paris, France; Department of Neuroscience, Icahn School of Medicine at Mount Sinai, New York, USA

## Abstract

Nicotinic acetylcholine receptors shape diverse brain functions, including reward, aversion, and pain, but dissecting their roles *in vivo* requires drug delivery with spatiotemporal precision that conventional pharmacology cannot provide. Here we establish *in vivo* photopharmacology of endogenous nicotinic receptors: two complementary highly-potent agonists, administered systemically in an inactive caged form, are photoreleased locally in target brain circuits of behaving wild-type mice. Despite irreversible photochemistry, photoactivation can be repeated over many trials on a minute timescale, for hours following a single injection. Local uncaging evokes circuit-specific dopamine release, drives place preference from the ventral tegmental area and aversion from the interpeduncular nucleus, and, in pain circuits, reveals a pronociceptive effect masked when the same agonists are given systemically. By controlling where and when nicotinic signaling acts, this approach exposes circuit-specific drug effects invisible to conventional pharmacology, and outlines a general strategy for on-demand control of endogenous receptors *in vivo*.

## Introduction

In the mammalian brain, acetylcholine (ACh) is synthesized in a small number of nuclei yet broadcasts widely, where it signals through muscarinic and nicotinic acetylcholine receptors (nAChRs). This anatomically restricted source and diffuse projection allows nicotinic signaling to shape a remarkably wide range of brain functions, from cognitive processes such as attention and memory, to reward encoding, aversion, anxiety, sleep, and pain (Picciotto et al., 2012; Bloem et al., 2014; Koukouli and Changeux, 2020). How a single neurotransmitter supports such functional diversity depends critically on where and when it is released: the effect of a given nicotinic signal is set by the neuronal pathway engaged, the timing of receptor activation and desensitization, and the mode of transmission - synaptic or volume - through which ACh reaches its targets (Sarter et al., 2009; Jing et al., 2020; Ananth et al., 2023). This picture is further complicated by the molecular diversity of nAChRs themselves: multiple subunit combinations (e.g., α4β2, α7, α3β4) assemble into pharmacologically and kinetically distinct receptor subtypes (Gotti et al., 2006).

Disentangling the functional roles of nicotinic neurotransmission requires methods for spatially and temporally controlled manipulation of nAChRs in the behaving mouse. Systemic nicotinic agonists activate receptor populations across the brain and the periphery simultaneously, so their net effects can obscure distinct or even opposing circuit-specific actions (Nguyen et al., 2021; Jehl et al., 2025), whereas local infusion remains limited by delivery and diffusion. Photopharmacology can, in principle, bridge this gap by separating drug delivery from receptor activation. There are three main photopharmacological tools: diffusible photoswitchable ligands, photoswitchable tethered ligands (PTLs), and caged compounds (Kramer et al., 2013; Hüll et al., 2018; Paoletti et al., 2019; Ellis-Davies, 2023; Colleoni et al., 2025).

Photoswitches have one significant advantage over caged compounds, in that the photochemical reaction is reversible. However, photoisomerization is incomplete, and in diffusible photoswitchable ligands both isomers typically retain activity at the receptor, with the thermally stable dark isomer usually being the most active one, so that a pharmacologically active species is already present in the tissue before any light is delivered. This is notably the case for photoswitchable nAChR agonists (Damijonaitis et al., 2015; Colleoni et al., 2025; Vangelatou et al., 2026), which, accordingly, have not been deployed in mice *in vivo*.

PTLs, unlike diffusible photoswitches, do reach near-binary control, but they pay for it with two requirements: genetic modification of the receptor and local delivery of the photoswitch, as illustrated with light-controllable nAChRs (Tochitsky et al., 2012; Durand-de Cuttoli et al., 2018; Sansalone et al., 2019). The local-delivery requirement is due to the maleimide anchor’s lability and broad reactivity (Paoletti et al., 2019; Ciscato et al., 2025b; Jehl et al., 2025).

In contrast, caged compounds use photochemical protecting groups attached to a crucial functionality, producing probes that are usually fully inert in the dark (Ellis-Davies, 2023), a property which diffusible photoswitches lack (Paoletti et al., 2019). Moreover, unlike tethered photoswitches, caged compounds allow photocontrol of endogenous receptors. Nicotine has been previously caged by two distinct approaches: Ruthenium–bipyridine-based RuBiNic, which efficiently releases nicotine under visible light (Filevich et al., 2010), and coumarin-based PA-Nic and its improved variant PANic2, which enabled subcellular mapping of nAChRs (Banala et al., 2018, 2024). Yet, none has been applied *in vivo* in the mammalian brain, in part because of unfavorable photochemistry and/or low agonist affinity.

Deploying caged compounds in living rodents is particularly challenging. To date, most caged neuroactive compounds validated in behaving mice have required local infusion through an opto-fluidic cannula. This is because the caged drug binds its target with low affinity and must be applied at high concentration, and/or because it does not cross the blood–brain barrier (Durand-de Cuttoli et al., 2020; Ma et al., 2023). Recently, a nitrobenzyl photochemical protecting group was shown to carry small-molecule opioid ligands across the blood–brain barrier, enabling systemic administration and local photorelease in behaving mice (McClain et al., 2023). However, whether the same approach could provide photocontrol of endogenous nAChRs, rapidly activating and desensitizing ligand-gated ion channels, remained unknown. In this study, we show that two nitrobenzyl-caged nicotinic agonists can be injected systemically into wild-type mice to control nAChR function and behavior with high spatiotemporal precision. The approach is compatible with standard neurophysiological techniques, including electrophysiology, fiber photometry and functional ultrasound imaging, suggesting that it could be broadly applicable to the photopharmacological interrogation of physiology and behavior.

## Results

### Design and synthesis of potent caged nicotinic agonists

We reasoned that pairing a blood–brain barrier-permeant neutral cage with a highly potent nicotinic agonist would enable effective photocontrol of brain nAChRs following systemic loading (**Fig. 1a**). To maximize the versatility of this approach, we selected two high-affinity nicotinic agonists with distinct pharmacological and pharmacokinetic profiles: epibatidine (EPI) and its clinically inspired analogue tebanicline (ABT594) (Spande et al., 1992; Sullivan et al., 1994; Bannon et al., 1998b; Lin et al., 2020). We caged both agonists by installing a near-UV sensitive nitroveratyl (NV) cage on the secondary amine required for receptor engagement (**Fig. 1b**), guided by structure–activity and structural data. Substituting the secondary amine proton of EPI with an ethyl or larger group sharply reduces affinity (Carroll, 2004), and installing a bulky NV group on the equivalent amine of ABT594 likewise yields a dark-inert agonist (Passlick et al., 2018). A high-resolution structure of the α7 nAChR bound to EPI shows this secondary amine buried deep in the orthosteric pocket, where NV substitution would clash sterically with Trp149, an essential binding-site residue (**Fig. 1c,d**) (Noviello et al., 2021). Because EPI is a high-affinity agonist principally of α4β2 nAChRs, we modeled caging in this heteromeric subtype. Docking of EPI and ABT594 into the α4β2 nAChR confirmed that both adopt similar, favorably scored poses resembling the α7-EPI complex, whereas NV-EPI and NV-ABT594 failed to reach productive orientations (**Fig. S1a-c**), precluding meaningful score comparison, consistent with steric occlusion of the agonist–receptor interaction by the NV group. NV-ABT594 was synthesized as previously described (Passlick et al., 2018), whereas NV-EPI was synthesized as described in **Fig. S2a-c**.

**Figure 1.**
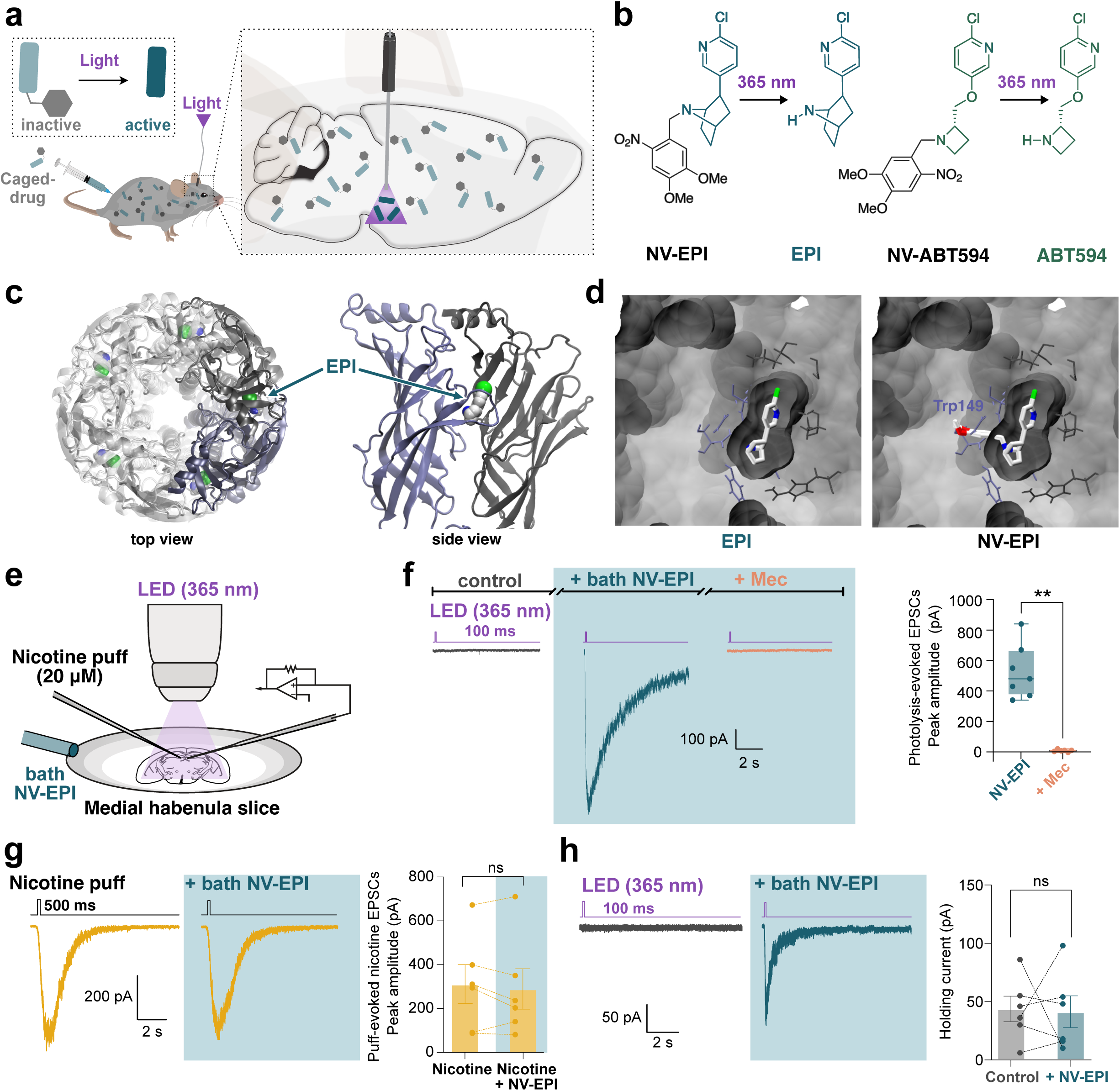
Design and validation of NV-caged nicotinic agonists for photopharmacological control of nAChRs. **a**, Schematics of *in vivo* photopharmacology using systemically administered caged drugs. Top insert, schematics of drug uncaging (drug in green, cage in black). Right insert, local drug photorelease in the brain using an implanted optical fiber. **b**, Chemical structures and photolysis reactions of NV-Epibatidine (NV-EPI) and NV-ABT594, showing 365-nm-triggered uncaging to yield EPI and ABT594, respectively. **c**, High-resolution structure of the homopentameric α7 nAChR bound to EPI (PDB 7KOQ, chloride atom in green), shown in top view (left) and side view (right). **d**, Left, close-up views of the agonist binding site showing EPI. Right, attachment of the NV cage on the secondary amine of EPI clashes sterically with the protein, notably residue Trp149. **e**, Schematic of whole-cell patch-clamp recording in medial habenula (MHb) brain slices, with full-field illumination for uncaging, bath application for NV-EPI and nicotine puff application. **f**, Left, a representative voltage-clamp recording (−60 mV) showing robust inward current evoked by a 100-ms pulse of 365-nm light in the presence of NV–EPI (100 nM). Currents were absent under control (no NV-EPI) condition and blocked by the non-selective, non-competitive nAChR antagonist mecamylamine (Mec, 10 µM). Right, average current amplitude (526 ± 67 pA, n = 7 cells) is abolished with Mec (8 ± 3 pA, n = 6, *p* = 0.001, Mann-Whitney test). **g**, Puff-evoked nicotine (20 µM) responses recorded before and after bath application of NV–EPI (100 nM). Bath-applied NV-EPI did not alter nicotine-evoked current amplitude (baseline, 309 ± 89 pA, n = 6; after NV-EPI, 287 ± 92 pA, n = 6; *p* = 0.43, Wilcoxon matched-pairs signed-rank test). **h**, A representative current produced upon 365-nm light stimulation (100 ms) in the absence or presence of NV–EPI (100 nM). NV-EPI did not alter the baseline holding current before illumination (43.2 ± 10.9 pA, n = 6; NV-EPI, 40.7 ± 13.6 pA, n = 6; *p* = 1.0, Wilcoxon matched-pairs signed-rank test).

### NV-EPI photoactivates endogenous nAChRs in brain slices

NV-ABT594 was validated previously on endogenous nAChRs in brain tissue (Passlick et al., 2018), therefore we focused our *ex vivo* characterization on the new compound, NV-EPI. We performed whole-cell patch-clamp recordings from medial habenula (MHb) neurons in acute brain slices (**Fig. 1e**), a region in which nAChRs are highly expressed (Gotti et al., 2006; Ciscato et al., 2025a). In voltage-clamp mode, brief pulses of 365-nm light delivered in the presence of bath-applied NV-EPI elicited large inward currents, indicating efficient photorelease of EPI, whereas identical illumination in the absence of caged compound produced no detectable response (**Fig. 1f**). These photocurrents were abolished by the nAChR antagonist mecamylamine, confirming that they are mediated by nAChR activation (**Fig. 1f**).

A critical requirement for any caged compound is that it remains pharmacologically inactive before photolysis, acting as neither an agonist nor an antagonist (Ellis-Davies, 2023). We tested the dark-state activity of NV-EPI in two complementary ways. First, inward currents evoked by repeated nicotine puffs were unchanged during bath application of NV-EPI, arguing against direct antagonism or receptor desensitization caused by residual agonist activity (**Fig. 1g**). Second, bath application of NV-EPI at the concentrations used experimentally did not alter the neuronal holding current, indicating no detectable tonic agonist activity (**Fig. 1h**). Together, these results show that NV-EPI is pharmacologically silent in the dark at the concentrations used in our experiments.

### Systemic NV-EPI and NV-ABT594 enable photocontrol of neuronal activity *in vivo*

Next, we recorded from ventral tegmental area (VTA) putative dopamine (DA) neurons in brain slices (**Fig. 2a-b**), a brain region central to nicotine reward (Maskos et al., 2005; Nguyen et al., 2021). Violet-light illumination in the presence of NV-EPI reliably evoked action potential firing (**Fig. 2c**), similarly to what was previously reported for NV-ABT594 (Passlick et al., 2018).

**Figure 2.**
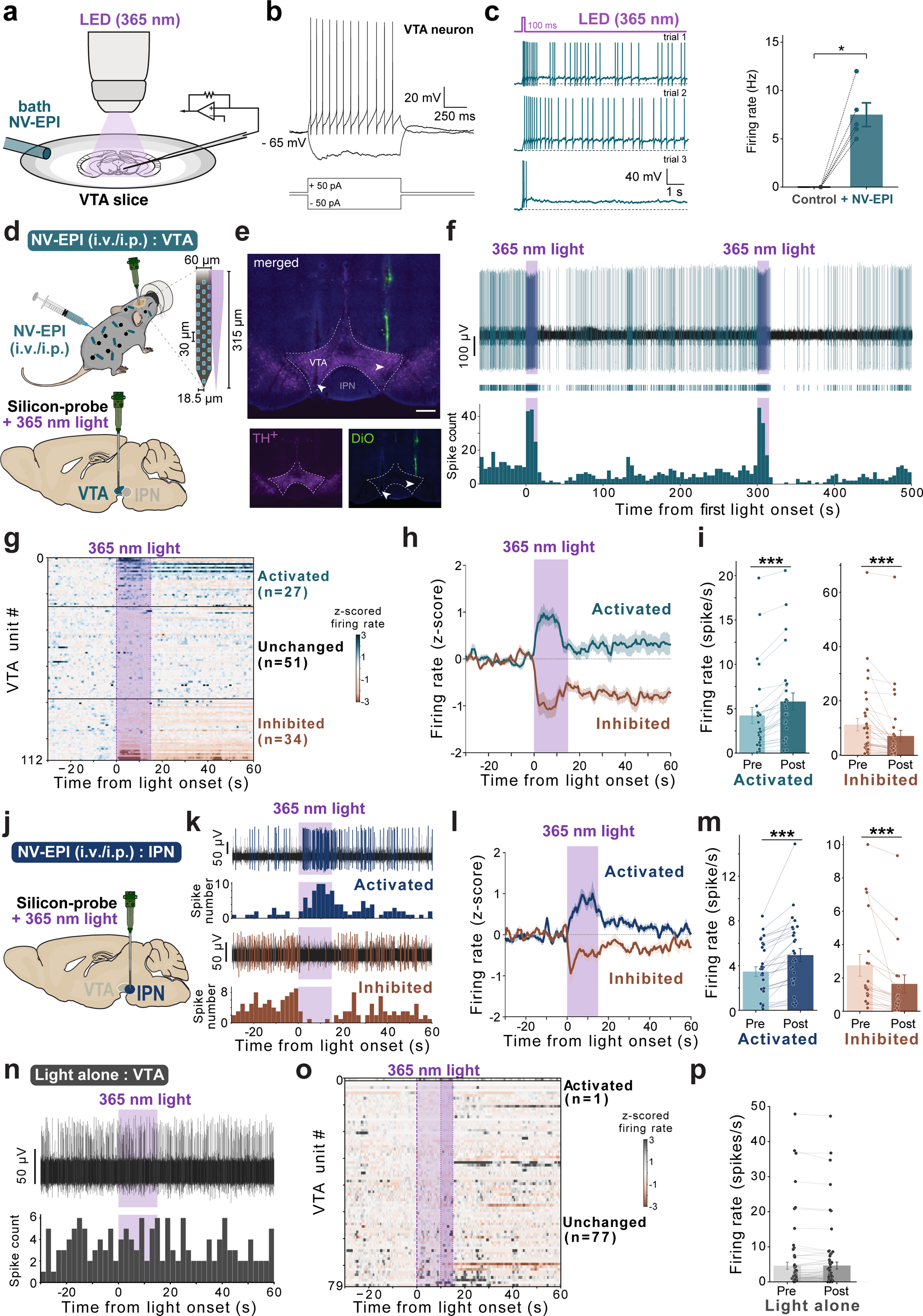
Systemic NV-EPI crosses the blood–brain barrier and enables photocontrol of VTA and IPN neurons. **a**, Schematic of whole-cell current-clamp recording in VTA brain slices with bath-applied NV-EPI (100 nM) and local 365-nm LED illumination. **b**, Representative current-clamp traces from a VTA neuron showing intrinsic firing properties in response to depolarizing (+50 pA) and hyperpolarizing (−50 pA) current injection, showing electrophysiological properties characteristic of putative dopaminergic neurons. **c**, Left, Representative traces of light-evoked action potential firing in VTA neurons across three trials following 365-nm illumination (100 ms) in the presence of bath-applied NV-EPI. Right, 365-nm illumination increased firing rate relative to control (control, 0.0 ± 0.0 Hz, n = 5; NV-EPI, 7.5 ± 1.2 spikes/s, n = 5; *p* = 0.03, one-sided Wilcoxon matched-pairs signed-rank test). Data are mean ± SEM. **d**, Schematic of the *in vivo* electrophysiology approach: NV-EPI administered systemically (i.v./i.p.) in anesthetized mice, with an optrode implanted for combined 365-nm illumination and extracellular recording, targeting VTA. **e**, Histological validation of probe placement: merged image (top) with TH^+^ immunostaining (bottom left) and DiO probe-tract labeling (bottom right); dashed lines outline VTA and IPN boundaries. Scale bar: 300 µm. **f**, Representative voltage trace (top) and corresponding spike-rate histogram (bottom, spikes/5 s) from a VTA neuron showing light-evoked firing re-evoked by two successive 365-nm light pulses delivered 300 s apart. **g**, Heatmap of mean z-scored firing rates for recorded VTA units (n = 112 units from 4 mice) aligned to the onset of the 15-s light stimulus and averaged across three trials. Units are sorted into activated (n = 27), unchanged (n = 51), and inhibited (n = 34) populations. **h**, Mean (± SEM) z-scored firing rates over time for activated and inhibited VTA populations, aligned to light onset. **i**, Firing rates (spikes/s) before and after light onset for individually activated (left) and inhibited (right) VTA units; bars show mean ± SEM. Activated units increased their firing rate (+1.6 spikes/s, 95% CI [1.11, 2.04], *p* < 0.001), whereas inhibited units decreased their firing rate (−4.19 spikes/s, 95% CI [−5.92, −2.47], *p* < 0.001). **j**, Schematic of optrode recording combined with 365-nm illumination targeted to the IPN, following systemic NV-EPI administration. **k**, Representative voltage traces and spike-rate histograms (spikes/2 s) from an activated (top) and an inhibited (bottom) IPN neuron, aligned to 365-nm light onset. **l**, Mean (± SEM) z-scored firing rate over time for activated (n = 30 units) and inhibited (n = 19 units) IPN populations, aligned to light onset. **m**, Firing rate (spikes/s) before and after light onset for individual activated (left) and inhibited (right) IPN units; bars show mean ± SEM. Activated units increased their firing rate (+1.48 spikes/s, 95% CI [0.98, 1.98], *p* < 0.001), whereas inhibited units decreased their firing rate (−1.12 spikes/s, 95% CI [−1.77, −0.46], *p* < 0.001). **n**, A representative voltage trace and spike-rate histogram from a VTA neuron during light delivery alone (no NV-EPI), showing no light-evoked change in firing. **o**, Heatmap of mean z-scored firing rate for recorded VTA units (n = 78 units from 3 mice), aligned to light onset. For each unit, firing rate was averaged across available trials (up to three trials per unit); light-stimulus duration was 10 or 15 s. Units are sorted as activated (n = 1) or unchanged (n = 77). **p**, Mean firing rate before and after light delivery for individual VTA units in light alone controls. The mean change was 0.059 spikes/s (90% CI, −0.131 to 0.249; 78 units, three mice and five recordings). The CI fell within the prespecified equivalence bounds of ±0.5 spikes/s (two one-sided tests, *p* < 0.001). Bars show mean ± SEM.

To test whether the caged agonists cross the blood–brain barrier and can act *in vivo*, we performed acute *in vivo* electrophysiological recordings in anesthetized mice. A silicon probe was targeted to the VTA and NV-EPI was administered systemically, either intravenously or intraperitoneally (i.v. or i.p., **Fig. 2d,e, S3a,b**). Individual VTA neurons exhibited clear responses to local, 15-s violet illumination, which could be re-evoked with a second light pulse 300 s later (**Fig. 2f**). Responses were similar after i.v. or i.p. injection, with comparable proportions of neurons activated and inhibited (**Fig. 2g-i, S3b,c**). This duality in the responses is consistent with previous reports showing that nicotine activates and inhibits distinct populations of VTA DA neurons *in vivo* (Nguyen et al., 2021; Le Borgne et al., 2025). Illumination as brief as 5s produced changes in firing frequency of similar amplitude (**Fig. S3d**). Systemic NV-ABT594 uncaged in the VTA gave similar results, with both activated and inhibited neurons (**Fig. S3e-i**), showing that both caged agonists cross the blood–brain barrier and can be photoactivated *in vivo*.

We repeated this experiment in the interpeduncular nucleus (IPN, **Fig. 2j, S4a**), a structure that mediates nicotine aversion and limits the drug’s rewarding effects (Wolfman et al., 2018; Wills et al., 2022; Mondoloni et al., 2023; Ciscato et al., 2025a; Jehl et al., 2025). As in the VTA, uncaging EPI (**Fig. 2k-m, S4b,c**) or ABT594 (**Fig. S4d-g**) produced both activated and inhibited neurons, consistent with the dual nature of nicotine responses reported previously (Mondoloni et al., 2023; Jehl et al., 2025). Light alone had no such effect on neuronal activity in either the VTA or IPN (**Fig. 2n-p, S4h-j**), confirming that the responses depend on photoreleased agonist rather than on illumination itself. Because i.v. and i.p. injection gave similar results (**Fig. S3b, S4b)**, we used the simpler intraperitoneal route in the following experiments.

### Photorelease is spatially restricted to the illuminated brain area

To investigate the spatial profile of brain activation associated with photorelease, we combined NV-EPI uncaging with functional ultrasound imaging (fUSi). fUSi monitors cerebral blood volume (CBV) at high spatiotemporal resolution (100 μm, 400 ms) and offers great sensitivity with which to investigate brain-wide responses to single transient events (Macé et al., 2011; Dizeux et al., 2019; Bergel et al., 2020). Mice were prepared with a cranial window for chronic fUS imaging and implanted unilaterally with an angled optical fiber mounted on a microdrive, allowing the fiber tip to be positioned to intersect coronal imaging plane encompassing both VTA hemispheres (**Fig. 3a,b**). Vascular responses were monitored during i.p. administration of NV-EPI and stimulation with 365-nm light (5, 10 and 15 s in randomized triplets; **Fig. 3c, Supplementary Video 1**). EPI photorelease evoked a robust vascular response in the stimulated VTA that was greater than in the contralateral VTA, indicating that activation was spatially restricted around the targeted fiber location (**Fig. 3d,e**). In addition, light alone produced little to no response, confirming that the vascular activation was due to EPI uncaging (**Fig. 3d,f**).

**Figure 3.**
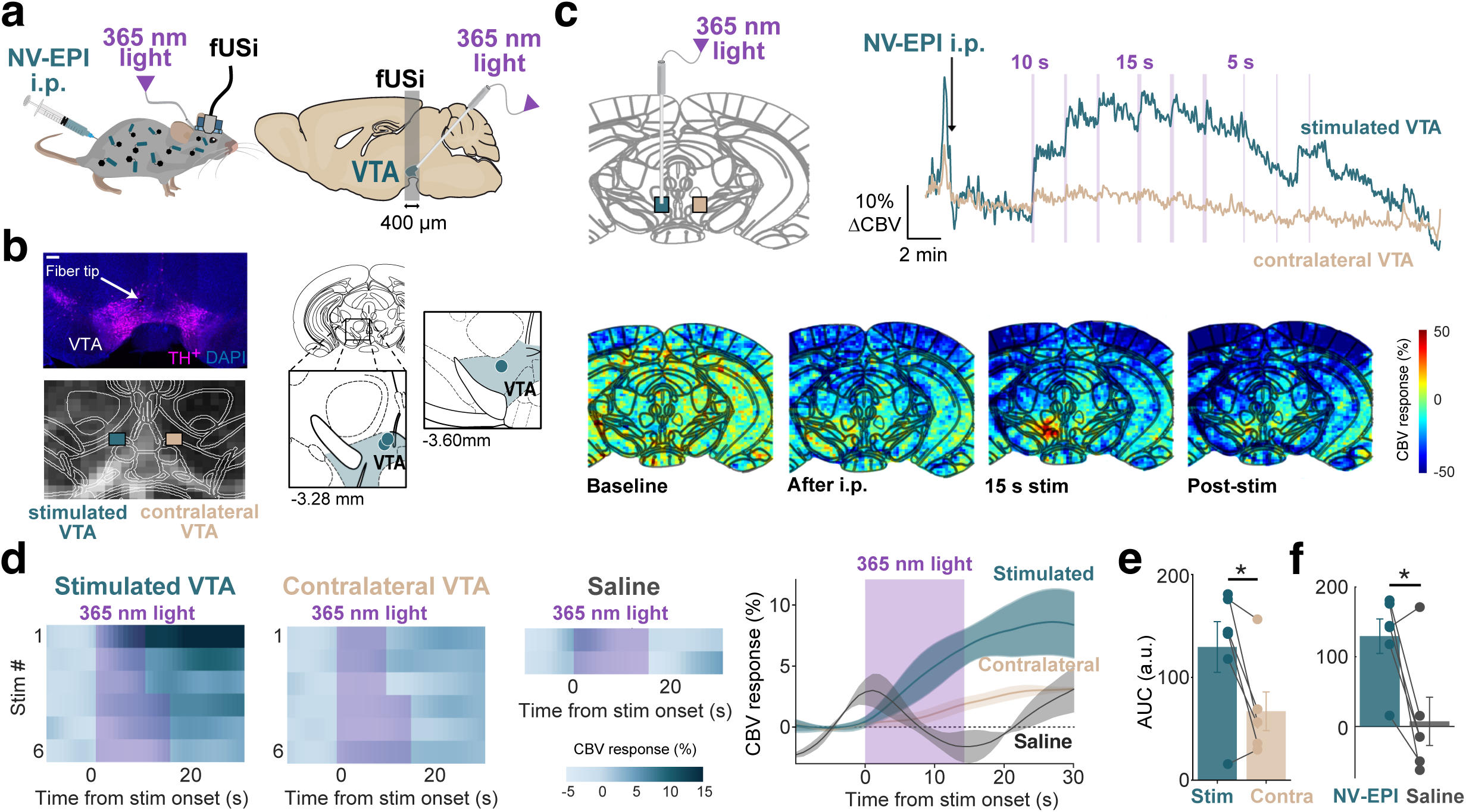
Spatially restricted photorelease measured by functional ultrasound imaging. **a**, Experimental design combining photopharmacology with fUSi. Mice were prepared with a cranial window for chronic fUS imaging and implanted unilaterally with an optical fiber targeting the VTA. During recording, cerebral blood volume data was acquired over a coronal plane encompassing both VTAs. **b**, Left, representative histological section showing the optical fiber tip relative to tyrosine hydroxylase-positive (TH^+^) neurons in the VTA and the corresponding fUS imaging plane (DAPI, purple; TH^+^, magenta). Scale bar: 200 µm. Square regions of interest (ROIs; 200 µm × 200 µm) were positioned at the estimated optical fiber tip in the stimulated VTA and at the corresponding contralateral site. Right, anatomical reconstruction of optical fiber tip positions in all three mice. **c**, Temporal profile of the vascular response in the stimulated and contralateral VTAs after i.p. administration of NV-EPI. Light stimuli of 5, 10 and 15 s were delivered in randomized triplets, with a 2-minute interval between successive stimuli. Representative fUS images (bottom) show a spatially restricted increase in cerebral blood volume in the stimulated VTA following photorelease of EPI. Shading indicates 365-nm light deliveries. **d**, Left, cerebral blood volume responses to the first six light stimuli in the stimulated VTA and corresponding contralateral ROI after NV-EPI injection, and of the first two light stimuli after saline administration (light alone condition). Right, mean responses from one representative mouse. **e**, Response area under the curve (AUC) in the stimulated and contralateral VTAs across six recording sessions from three mice. The response was greater in the stimulated VTA than in the contralateral ROI (mean paired difference = 62.5 AUC units, bootstrap 95% CI 23.2 to 99.2; paired two-sided Student’s t-test, *p* = 0.03). **f**, Response AUC after NV-EPI or saline administration across six recording sessions from three mice. Photorelease of EPI produced a greater vascular response than light delivery after saline administration (mean paired difference = 122.1 AUC units, bootstrap 95% CI 50.6 to 184.2; paired two-sided Student’s t-test, *p* = 0.02).

### Rapid, repeated and graded uncaging of nicotinic agonists *in vivo*

In order to assess further the temporal precision, repeatability and robustness of our method, we combined local uncaging with fiber photometry. We expressed the fluorescent DA sensor GRAB_DA2m_ (Sun et al., 2020) in the nucleus accumbens (NAc) and implanted optical fibers in this region as well as in the ipsilateral VTA, its principal dopaminergic input (**Fig. 4a, S5a**). NV-EPI or NV-ABT594 were injected i.p., and NAc DA dynamics served as a readout of VTA DA neuron photostimulation. Following a single i.p. injection of NV-EPI, a 15 s 365-nm light pulse to the VTA produced large, time-locked DA signals in the NAc, unlike in saline-injected, light-alone controls (**Fig. 4b**). The signal decayed within seconds of light offset. For NV-ABT594, 1 s violet pulses were sufficient to elicit large DA responses that persisted for many seconds after light offset, with no response in light-alone controls (**Fig. 4c**).

**Figure 4.**
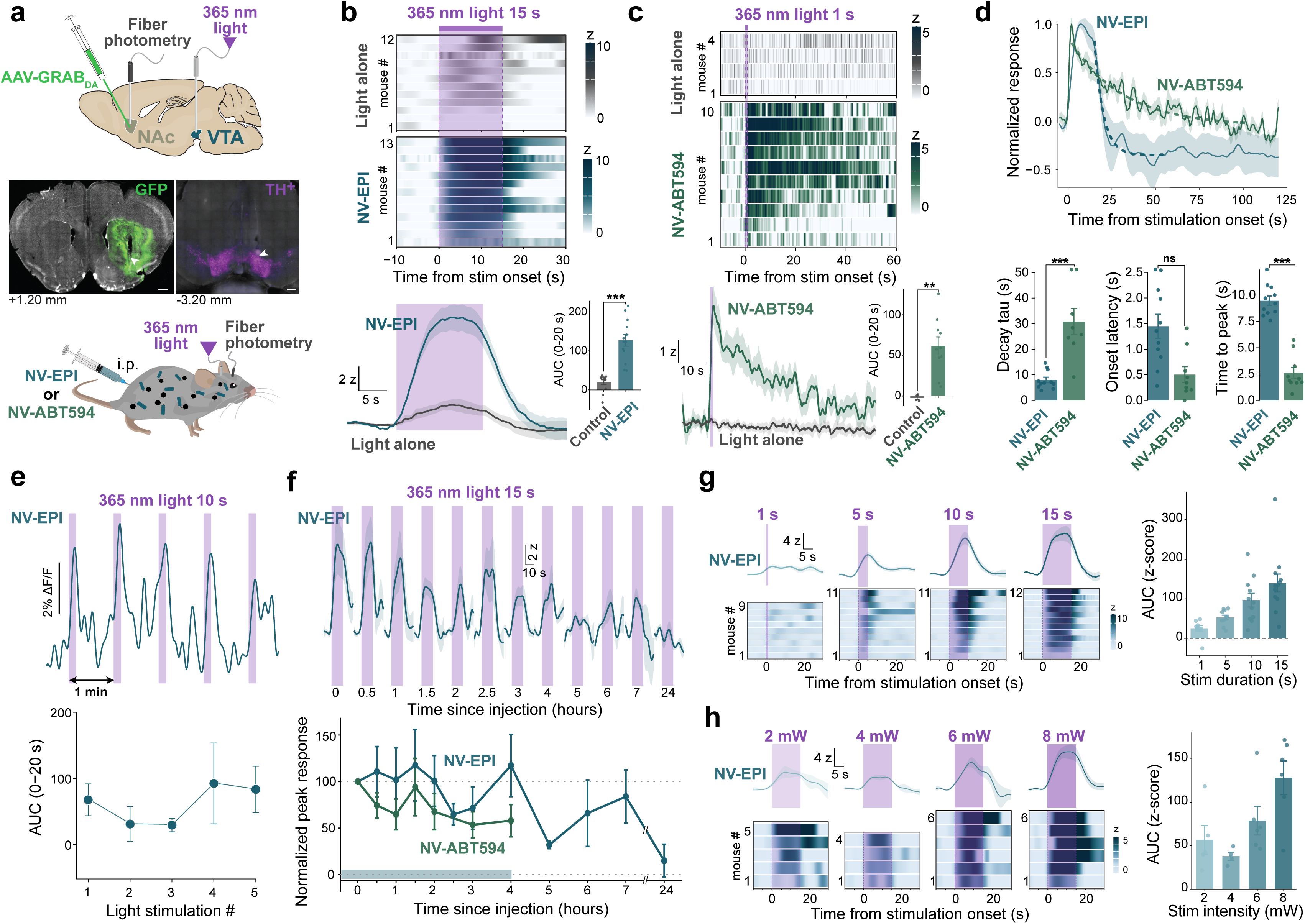
EPI and ABT594 uncaging drives rapid, repeatable, and graded dopamine release in the NAc. **a**, Top, schematic of the experimental approach: the fluorescent dopamine sensor GRAB-DA expressed in the nucleus accumbens (NAc), with fiber photometry in NAc, and a separate optical fiber delivering 365-nm light to the ipsilateral VTA. Middle, representative histology showing GRAB-DA (GFP^+^) expression in NAc (left) and TH^+^ immunostaining marking VTA dopaminergic neurons/fiber placement (right); scale bars: 500 µm and 300 µm, respectively. Coronal levels relative to bregma indicated. Bottom, fiber photometry recording in wild-type mice receiving systemic (i.p.) NV-EPI (200 µg/kg) or NV-ABT594 (4 mg/kg). **b**, Top, raster plot showing changes in NAc DA release (83 ms bin size) upon light stimulation (365-nm, 15 s, 8 mW) in saline control (n = 12 mice) and after i.p. injection with NV-EPI (n = 13 mice). Each row represents mean of mice averaged over three trials. Middle, average time course of the response. Right, response magnitude quantified as area under the curve (AUC). Uncaging of NV-EPI produced significantly larger dopamine responses than light alone (linear mixed-effects model, *F*(1,151.35) = 45.7, *p* < 0.0001; NV-EPI: 127 ± 15 z·s, n = 13 mice; light only: 19.4 ± 6.3 z·s, n = 12 mice; mean ± SEM of per-mouse means). **c**, As in **b**, for NV-ABT594 uncaging using a 1 s violet-light pulse. Uncaging of NV-ABT594 (n = 10 mice) produced significantly larger responses than light alone (n = 4 mice), measured by AUC over 0-20 s (NV-ABT594: 61.6 ± 11.0 z·s; light alone: −2.3 ± 1.8 z·s; LMM, *F*(1,11.19) = 13.47, *p* = 0.004). **d**, Time course of the normalized response to light for NV-EPI (15 s light pulse; n = 11 mice) and NV-ABT594 (1 s light pulse; n = 10 mice). Decay kinetics were fitted with a mono-exponential function. NV-EPI showed a shorter decay time constant than NV-ABT594 (τ: 7.97 ± 1.08 versus 30.8 ± 5.1 s; LMM, *F*(1,37) = 53.33, *p* < 0.0001) and a longer time to peak (9.5 ± 0.4 versus 2.6 ± 0.5 s; *F*(1,60) = 30.50, *p* < 0.0001). NV-EPI showed a trend toward a longer onset latency (1.45 ± 0.23 versus 0.50 ± 0.15 s; *F*(1,55.76) = 3.70, *p* = 0.059). Values are mean ± SEM of per-mouse measurements. **e**, Rapid repeatability of NV-EPI uncaging. Top, representative fluorescent trace (ΔF/F) from one mouse, showing repeated photoactivation (10-s light pulses, 1 minute apart) without rundown after a single IP injection of NV-EPI. Each stimulation epoch (violet shading) produces a robust DA transient. Bottom, AUC of the z-scored DA response (0-20 s window) plotted against stimulation number (1-5; mean ± SEM). Response amplitude was stable across all five violet stimulations (LMM: *F*(4,12) = 0.72, *p* = 0.59; linear trend *p* = 0.37). **f**, Repeatability of EPI and ABT594 uncaging over hours. Top, averaged traces (mean ± SEM) aligned to 15s violet stimulations delivered 0-7 hours and 24 hours after NV-EPI injection (200 µg/kg, i.p.; n = 4-8 mice per time point; exact n values are provided in the source data). Bottom, time course of peak z-scored DA responses following EPI or ABT594 uncaging. Peak responses were normalized within each mouse to the first stimulation after injection (100%). Filled circles with error bars show the mean ± SEM of normalized per-mouse values at each time point (NV-EPI, n = 4-8 mice per time point; NV-ABT594, n = 8-10 mice per time point). EPI could be reliably uncaged for at least 4 hours after injection, with peak responses remaining significantly greater than zero at 4 hours (6.66 ± 1.64 z-score, Holm-adjusted one-sided one-sample *p* = 0.026). Peak responses became less consistent at later time points, with no detectable positive uncaging response remaining at 24 hours (0.48 ± 0.76 z-score, Holm-adjusted one-sided one-sample *p* = 0.287). ABT594 could be reliably uncaged for up to 4 hours after injection, with peak responses remaining significantly greater than zero at every measured time point from 0 to 4 hours (Holm-adjusted one-sided one-sample tests, all *p* ≤ 0.009, 3.33 ± 0.79 z-score, *p* = 0.007). Together, these results support a reliable 0–4 hours photostimulation window for both compounds, indicated by green shading. **g**, Illumination-duration dependence of NV-EPI-evoked responses. Left: mean traces aligned to 365-nm-light stimulation of different durations (1, 5, 10 or 15 s; shaded area). Traces show mean ± SEM. Right: integrated dopamine responses (AUC over 0-20 s) increased with stimulation duration. Group means (bars, mean ± SEM across mice) and individual mouse means (dots) are shown for each duration (n = 9-12 mice per duration; 13 mice overall). A linear mixed model revealed a significant main effect of duration (*F*(3,28.78) = 10.35, *p* < 0.0001) and a positive linear relationship between duration and AUC (slope = 8.39 ± 1.45 z·s per second, t(32.68) = 5.78, *p* < 0.0001). **h**, Illumination-intensity dependence of NV-EPI-evoked responses. Left: mean traces and per-mouse heatmaps aligned to 365-nm-light stimulation at 2, 4, 6, and 8 mW (25-100 % maximum power; n = 4-6 mice per intensity). Right: AUC increased with illumination intensity overall (linear mixed-effects model with mouse as a random effect: *F*(3,17) = 5.34, *p* = 0.009), with a significant positive linear relationship between power and AUC (slope = 1.02 ± 0.31 AUC units per percentage-point increase in power, t(19) = 3.27, *p* = 0.004)

Both caged compounds evoked DA responses of peak amplitude comparable to that produced by a systemic dose of nicotine known to be reinforcing (0.5 mg/kg) (Jehl et al., 2025), but with dramatically sharper kinetics (**Fig. S5b-e**): faster time to peak and much shorter duration (a few seconds vs many minutes in both cases). Also, the two compounds differed sharply from each other: NV-EPI showed slower onset and time to peak but a much shorter decay constant than NV-ABT594 (**Fig. 4d**). Consequently, photoactivation could be repeated every minute without noticeable rundown for NV-EPI (**Fig. 4e**) but less so for NV-ABT594 (**Fig. S6a**).

Next, we asked how long a single injection supports reliable uncaging. Mice were stimulated every 30-60 minutes for up to 8 hours and again at 24 hours post-injection. DA responses remained significantly above baseline up to 4 hours for both, while none was detectable at 24 hours for NV-EPI (**Fig. 4f, S6b**). Thus, a single systemic injection supports repeated, temporally precise photostimulation over several hours.

A key advantage of optical control is the ability to tune responses by varying light duration or intensity (Paoletti et al., 2019; McClain et al., 2023). Isolated light pulses of increasing duration (1, 5, 10, 15 s) produced responses that grew monotonically with duration for both agonists (**Fig. 4g, S7**). Varying intensity (25-100% max power, 2-8 mW) at fixed duration (15 s) produced similar graded increases (**Fig. 4h**). Thus, the amplitude of DA release evoked by a single systemic injection can be scaled precisely across a dynamic range.

### Caging strongly attenuates the systemic locomotor effects of EPI

To assess the extent to which caging suppresses the behavioral activity of EPI, mice received i.p. injections of EPI or NV-EPI at escalating doses and were placed in the open field 5 minutes later for 30 minutes of video-tracked locomotion (**Fig. 5a**). EPI produced a robust, dose-dependent suppression of locomotor activity, approaching complete immobility at the highest doses tested (10 μg/kg, **Fig. 5b**), in agreement with previous reports (Bannon et al., 1995; Kesingland et al., 2000). In contrast, NV-EPI was essentially inactive across the same dose range, producing no detectable change in locomotion until the highest dose tested (200 µg/kg), with the dose–response curve shifted approximately 50-fold to the right (**Fig. 5b**). Moreover, at 200 µg/kg NV-EPI, locomotion fully recovered within one hour after injection (**Fig. 5c**), validating the use of this dose and a 1-hour pre-injection window for subsequent photopharmacology experiments *in vivo*.

**Figure 5.**
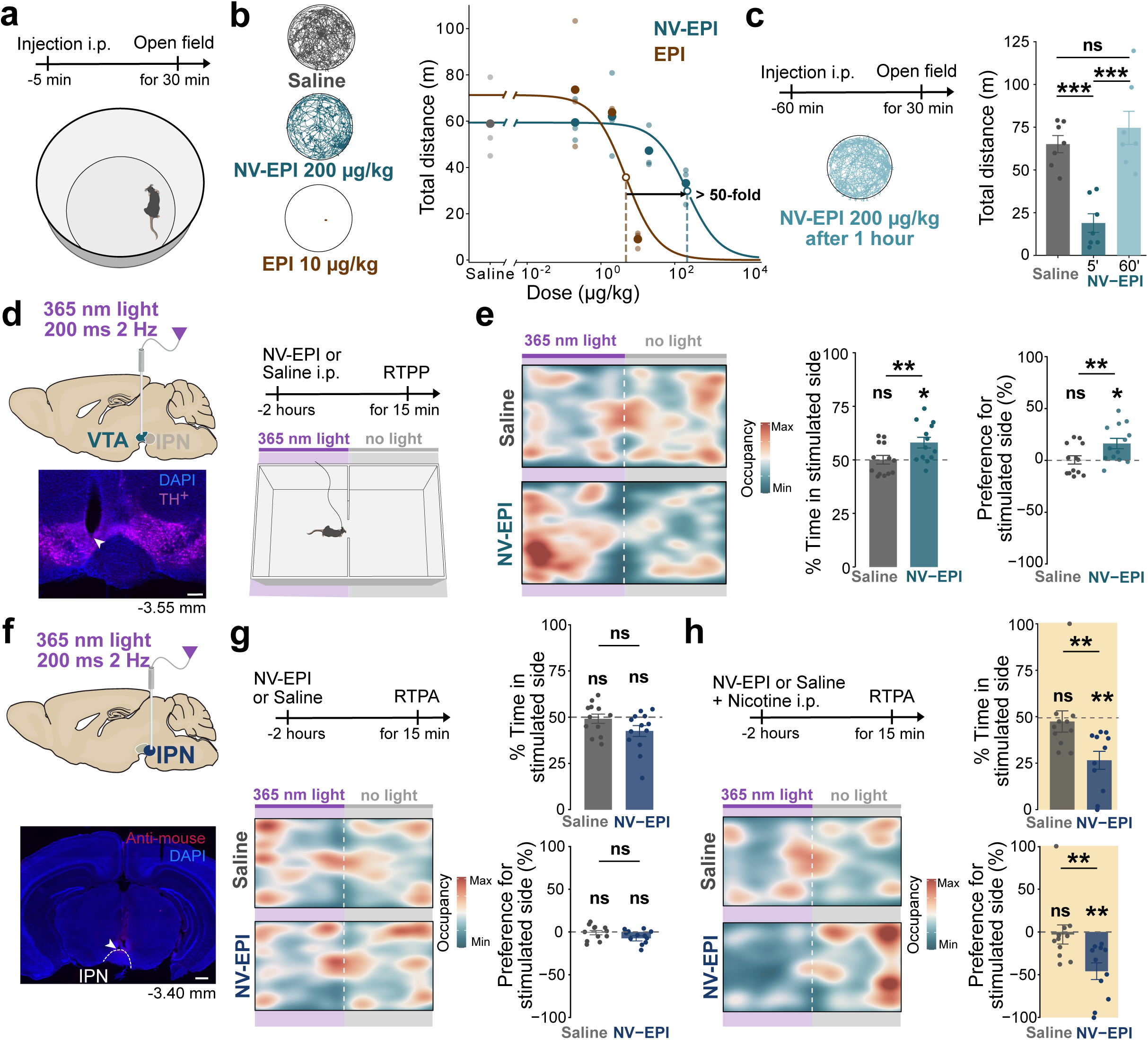
Local photorelease of EPI drives circuit-specific, bidirectional place preference/aversion. **a**, Schematic of the open-field locomotor assay: mice received i.p. injection of EPI or NV-EPI at escalating doses, 5 minutes before 30 minutes of video-tracked locomotion in an open-field arena. **b**, Left, representative locomotor tracks for saline, NV-EPI (200 µg/kg) or EPI (10 µg/kg). Right, dose-response curve for total distance traveled as a function of injected dose (µg/kg, log scale) for NV-EPI and EPI, showing a ∼50-fold right-shift for the EC_50_ of the caged drug (EPI EC_50_ = 4.8; NV-EPI EC_50_ = 217). n = 3 mice per dose. **c**, Left, schematic of the recovery assay. Mice received NV-EPI (200 µg/kg, i.p.) and were tested in the open field 60 minutes post-injection. Right, NV-EPI reduced total distance travelled at 5 min, but locomotion recovered to saline levels by 60 min. One-way ANOVA with Bonferroni-adjusted post hoc comparisons: *F*(2,18) = 18.07, *p* = 4.97 × 10⁻⁵; saline vs 5 min, *p* < 0.0001; saline vs 60 min, *p* = 1.0000; 5 vs 60 min, *p* < 0.0001. n = 7 mice per group. **d**, Schematic of the real-time place preference (RTPP) approach, with 365-nm light (200 ms, 2 Hz, 8 mW) delivered to the VTA upon chamber entry, following NV-EPI or saline injection (i.p., 2 hours before testing) and 15-minute testing. Bottom-left, representative histology showing fiber placement in the VTA (TH^+^ immunostaining, magenta; DAPI, blue); scale bar: 200 µm. **e,** RTPP results for VTA stimulation. Left, representative occupancy heatmaps for saline (top) and NV-EPI (bottom) groups across the light-paired and non-paired chambers. Middle, percent time spent in the stimulated side. NV-EPI mice spent more time in the stimulated side than saline-treated mice (NV-EPI, 58.0 ± 2.5%; saline, 50.0 ± 2.0%; Wilcoxon rank-sum test, *W* = 129, *p* = 0.02; *n* = 13 mice per group). One-sample tests against chance occupancy showed that NV-EPI mice were significantly above 50% (one-sample Wilcoxon signed-rank test, *W* = 83, *p* = 0.01), whereas saline-treated mice did not differ from chance (*W* = 51, *p* = 1.0). Right, preference score toward the stimulated side, showing increased preference in NV-EPI mice relative to saline-treated mice (NV-EPI, 16.3 ± 4.9%; saline, 0.6 ± 4.0%; Wilcoxon rank-sum test, *W* = 128, *p* = 0.025). **f**, Top, schematic of the real-time place aversion (RTPA) approach, with 365-nm light delivered to the IPN following NV-EPI or saline injection. Bottom, representative histology showing fiber placement in the IPN (anti-mouse antibody, red; DAPI, blue); scale bar: 500 µm. **g**, RTPA results for IPN stimulation in naive (non-nicotine-primed) mice. Left, representative occupancy heatmaps. Top-right: percent time spent in the stimulated side. NV-EPI mice showed a weak aversion-like tendency but did not differ significantly from saline-treated mice (saline, 49.2 ± 2.5%; NV-EPI, 42.6 ± 3.0%; unpaired *t*-test, *t*(23) = 1.70, *p* = 0.103; saline, *n* = 12; NV-EPI, *n* = 13). One-sample tests against chance occupancy showed that neither group differed significantly from 50% after correction (saline: one-sample *t*-test, *t*(11) = −0.34, *p* = 1.0; NV-EPI: one-sample *t*-test, *t*(12) = −2.52, *p* = 0.05). Bottom-right, preference score toward the stimulated side was −0.8 ± 2.5% for saline-treated mice and −7.4 ± 3.0% for NV-EPI mice. This difference was not statistically significant (unpaired *t*-test, *t*(23) = 1.70, *p* = 0.1; saline, *n* = 12; NV-EPI, *n* = 13). One-sample tests against neutrality (0%) showed that neither group differed significantly from zero after correction (saline: one-sample *t*-test, *t*(11) = −0.34, *p* = 1.0; NV-EPI: one-sample *t*-test, *t*(12) = −2.52, *p* = 0.05), consistent with the weak, non-significant aversion-like tendency observed in the stimulated-side occupancy analysis. **h**, RTPA results for IPN stimulation in nicotine-primed mice (nicotine i.p. co-administered, −2 h). Left, representative occupancy heatmaps. Top-right: NV-EPI mice spent 26.5 ± 4.9% of time in the stimulated chamber, compared with 47.4 ± 5.7% for saline-treated mice (Wilcoxon rank-sum test, *W* = 102, *p* = 0.007; *n* = 11 mice per group). NV-EPI mice also spent significantly less time in the stimulated chamber than expected by chance (50%; Wilcoxon signed-rank test, *W* = 0, *p* = 0.007), whereas saline-treated mice did not differ from chance (*W* = 14, *p* = 0.2). Bottom-right, preference score toward the stimulated side was −45.9 ± 9.8% for NV-EPI mice and −3.0 ± 11.2% for saline-treated mice. NV-EPI mice showed a significantly lower preference score than saline-treated mice (Wilcoxon rank-sum test, *W* = 102, *p* = 0.007; *n* = 11 mice per group). One-sample tests against neutrality (0%) showed that NV-EPI mice were significantly below zero (Wilcoxon signed-rank test, *W* = 0, *p* = 0.007), whereas saline-treated mice did not differ from neutrality (*W* = 18, *p* = 0.4), consistent with an aversion-like response following nicotine priming.

### Uncaging in the VTA drives place preference

To test whether EPI uncaging could produce a behaviorally relevant output in freely moving animals, we first used a real-time place preference (RTPP) paradigm in which light stimulations were delivered in the VTA upon entry into one chamber of a two-chamber arena (**Fig. 5d, S8a**). Uncaging of EPI caused a preference for the stimulation-paired chamber to develop, whereas saline controls remained neutral (**Fig. 5e**). This is consistent with prior reports showing intra-VTA nicotine infusion drives reinforcement (Maskos et al., 2005). Time-resolved analysis revealed the clearest between-group separation during the 5–15-minute interval, with no significant difference in the first 5 minutes and only a trend in the final bin (**Fig. S8b**), suggesting that preference developed progressively over the course of the session. These results demonstrate that spatially restricted uncaging in the VTA is sufficient to drive reward-like behavior in freely moving mice.

### Uncaging in the IPN drives place aversion

To assess the spatial specificity of the method, and probe its capacity to produce opposing behavioral outputs from neighboring but functionally distinct circuits, we targeted the IPN, a ventral midbrain structure lying close to the VTA (**Fig. 5f, S8a**). In naive mice, EPI uncaging produced transient place aversion, significant only during the first 5 minutes (**Fig. 5g**, **S8c**). This aversion was enhanced strongly by nicotine priming: mice that received a prior systemic nicotine injection showed markedly reduced occupancy of the stimulation-paired chamber, an effect that was most pronounced during the first 5 minutes (**Fig. 5h, S8d**), consistent with prior work showing that nicotine priming is required for optogenetic stimulation of IPN neurons to become aversive (Morton et al., 2018). Locomotor activity did not differ between groups (**Fig. S8e**), arguing against a gross motor explanation for the observed chamber bias.

### Caging abolishes off-target locomotor effects in NV-ABT594

The behavioral inertness of NV-ABT594 was tested, as for NV-EPI, using mouse locomotion as a readout. ABT594 profoundly reduced locomotion at its effective dose, as previously reported (Kesingland et al., 2000; Meyer et al., 2006), while NV-ABT594 had no significant effect on locomotion across a wide dose range (**Fig. 6a**). Only at a dose more than 300-fold higher than the parent compound’s effective dose did we observe a small, yet significant suppression. These results confirm a substantial widening of the behavioral safety window upon caging ABT594.

**Figure 6.**
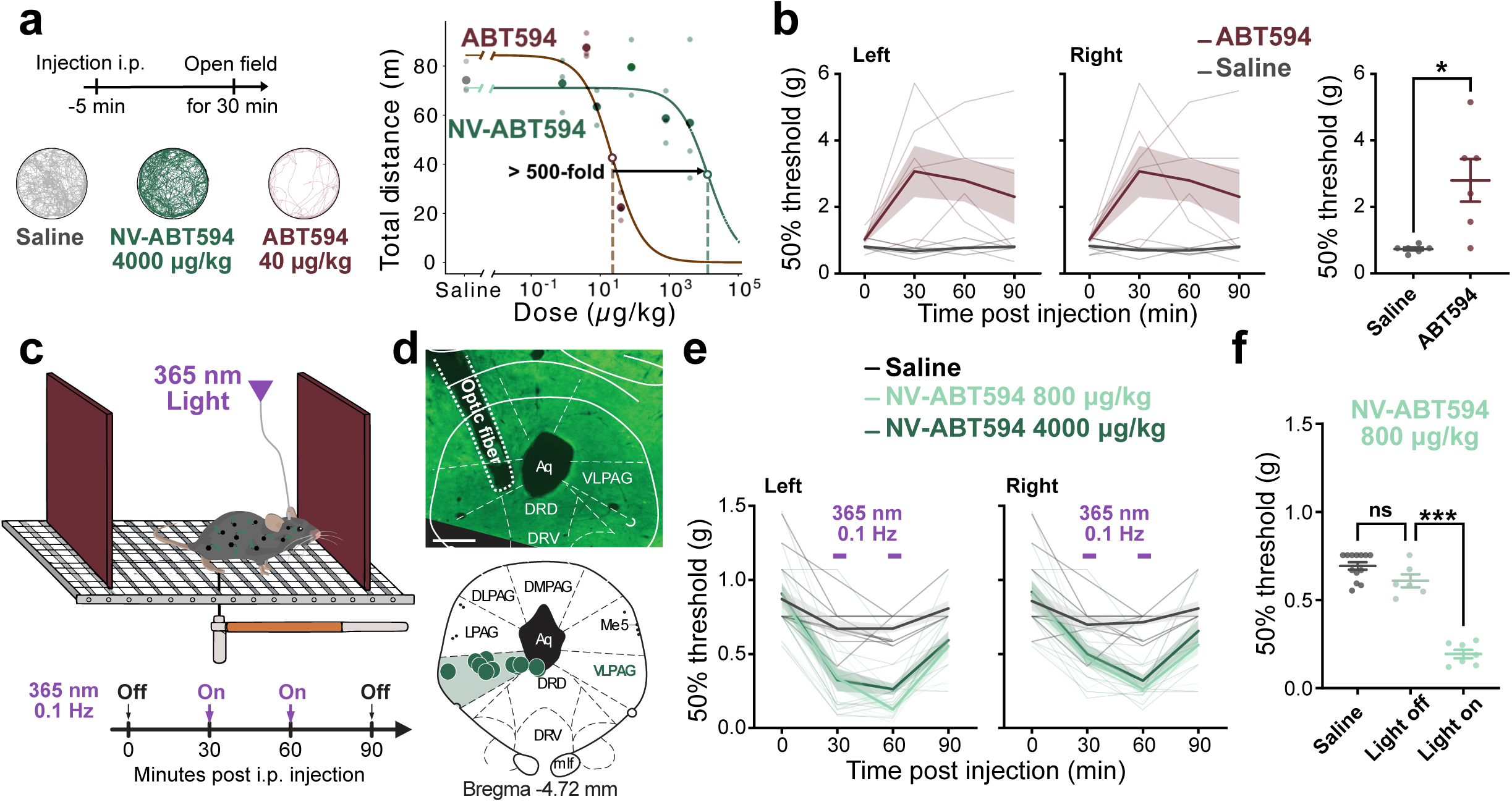
Systemic and locally photoreleased ABT594 exert opposing effects on mechanical sensitivity **a**, Left, schematic of the open-field locomotor assay (i.p. injection, 30-minutes video-tracked locomotion 5 minutes later). Representative locomotor tracks for saline, NV-ABT594 (4000 µg/kg), and ABT594 (40 µg/kg) shown below. Right, dose–response curves for total distance traveled as a function of injected dose (µg/kg, log scale) for ABT594 and NV-ABT594; the NV-ABT594 curve is shifted >300-fold rightward relative to ABT594 (ABT594 EC_50_ = 23; NV-ABT594 EC_50_ = 11250. n = 3 mice per dose. **b**, Systemic ABT594 increases mechanical withdrawal thresholds. Left and middle, Time course of the 50% mechanical withdrawal threshold (g; von Frey test) in the left and right hind paws following saline or ABT594 injection. Thin lines represent individual mice; thick lines and shaded areas indicate the group mean ± SEM. Right, Mean mechanical withdrawal threshold 60 minutes after injection. ABT594 significantly increased the withdrawal threshold compared with saline (*p* = 0.03, Wilcoxon signed-rank test; n = 6 mice). Each dot represents an individual mouse; bars indicate the mean ± SEM. **c**, Schematic of the experimental paradigm used to assess mechanical sensitivity in mice. Animals were placed on an elevated grid, and mechanical withdrawal thresholds were measured using calibrated von Frey filaments. NV-ABT594 or saline was administered i.p., and mechanical thresholds were assessed at baseline and at the indicated time points post-injection. 365-nm light (0.1 Hz) was applied during the indicated epochs. **d**, Validation of fiber implantation sites. Top, representative image showing optical fiber placement in the ventrolateral periaqueductal gray (vlPAG). Bottom, schematic representation of implantation coordinates across animals plotted on the mouse brain atlas (Bregma −4.72 mm anteroposterior axis), highlighting the targeted brain regions. Scale bar: 500 µm. **e**, Mechanical withdrawal thresholds for the left (left panel) and right (right panel) hind paws at different time points following i.p. administration of NV-ABT594 (0.8 mg/kg or 4 mg/kg) or saline. 365-nm light stimulation was applied at 30 and 60 minutes post-injection, as indicated by a purple line. Local photorelease of NV-ABT594 in the vlPAG produced the opposite behavioral effect from systemic ABT594. During light stimulation, 50% von Frey withdrawal thresholds decreased at 30 and 60 min, indicating increased mechanical sensitivity. Thresholds returned toward baseline at 30 minutes after light stimulation was switched off (90 minutes after injection). This light-dependent decrease was observed at both 4 mg/kg and 0.8 mg/kg NV-ABT594 and was present in both left and right paws. Saline-treated animals showed no comparable light-dependent effect. Thin lines represent individual animals; thick lines show the group mean ± SEM. **f**, Quantification of the mean mechanical withdrawal threshold at 60 minutes post-injection under saline, 365-nm light-off, and 365-nm light-on conditions. NV-ABT594 significantly decreased withdrawal threshold only during light stimulation, indicating that the hypersensitivity was triggered by local photorelease (Mixed effects analysis followed by Sidak’s multiple comparisons test, saline vs. light off, adjusted *p* = 0.14; light off vs. light on, adjusted *p* < 0.001). Each dot represents an individual animal; bars indicate mean ± SEM.

### Photoactivation reveals local pronociceptive action in the periaqueductal gray

Nicotinic agonists have long been explored as potent non-opioid analgesics, and EPI and ABT594 produce robust antinociception in preclinical models (Bannon et al., 1998b, 1998a). Clinical development of this class has been limited by adverse effects and by an incomplete understanding of the circuit-level sites at which nicotinic agonists produce their analgesic effects (Meyer, 2006). nAChRs are broadly distributed across spinal and supraspinal nodes of the pain pathway, several of which can contribute to pain modulation (Cheng et al., 2026), yet, where along these circuits nicotinic analgesia arises remains unclear. Consistent with previous reports of systemic nicotinic analgesia, we found that i.p. ABT594 increased paw withdrawal thresholds relative to saline (**Fig. 6b**), indicating mechanical hyposensitivity.

So, we asked whether the periaqueductal gray (PAG), a major node of descending pain control that expresses nAChRs (Nakamura and Jang, 2010; Umana et al., 2013; Sullere et al., 2023; van Strien and Hollmann, 2025), contributes to this modulation. Optical fibers were implanted unilaterally into the ventrolateral PAG (vlPAG), NV-ABT594 was injected i.p., and mechanical nociception was probed, as with ABT594, using von Frey filaments (**Fig. 6c,d**). We used NV-ABT594 for these experiments, as it has more sustained response after photorelease than NV-EPI (**Fig. 4d**). Surprisingly, local photorelease of NV-ABT594 in the vlPAG produced the opposite behavioral effect to systemic dosing (**Fig. 6b**): withdrawal thresholds decreased for left and right paws during illumination and recovered after light offset, with no effect of light alone (**Fig. 6e,f**). Focal activation of nAChRs in the vlPAG thus reveals a pronociceptive action masked during systemic exposure of ABT594.

## Discussion

We have developed two caged nicotinic agonists that, after systemic administration in an inactive form, can be photoreleased in the brain of freely moving mice through an implanted optical fiber. Penetrance of drugs in the brain remains a central obstacle in neuropharmacology; we addressed it by selecting a neutral NV cage, following evidence that neutral nitrobenzyl-type cages, unlike charged variants, cross the blood–brain barrier in mice (McClain et al., 2023; Li et al., 2026). Furthermore, by caging two high-affinity agonists, we could work at low systemic doses. Together, these properties enabled a single systemic injection support local and repeated photorelease in behaving mice.

One crucial advantage of our method is the ability to photocontrol endogenous receptors, in their normal subtype composition, subcellular distribution, and density, an advantage for questions where receptor fidelity is itself the object of study. The absence of any genetic requirement also removes the expression step that gates optogenetic, chemogenetic, and PTL methods (Mondoloni et al., 2019; Paoletti et al., 2019), making it applicable in wild-type animals and, in principle, across species and at any developmental stage at which light delivery is feasible. Furthermore, because uncaging delivers the agonist without fluid flow or tissue displacement, unlike pressure-driven infusion, it is readily combined with simultaneous *in vivo* electrophysiological recordings and fUSi.

The two caged agonists characterized here offer complementary temporal and physiological profiles, i.e. brief, rapidly repeatable responses with NV-EPI, versus more sustained responses with NV-ABT594, that can be matched to the timescale of the biological question. Because nAChRs desensitize rapidly (Quick and Lester, 2002), the response they produce depends on how quickly agonist concentration rises: slow delivery, as in local infusion or systemic injection, may drive receptors preferentially toward desensitization (Matta et al., 2007), whereas rapid photorelease presents the agonist as a near-step increase and engages the activated state of the receptors preferentially, at least on the rising phase of the response. Although uncaging is irreversible intrinsically, photostimulation could be repeated multiple times with both caged compounds; for NV-EPI, responses could be re-evoked at minute-scale inter-stimulus intervals with no detectable rundown, likely due to diffusional replenishment of intact caged compound beneath the fiber tip.

The spatial resolution of photostimulation was supported by two complementary lines of evidence. fUSi shows that photorelease evoked hemodynamic activity confined to the tissue immediately beneath the fiber tip, providing a direct physical readout of the volume recruited. This anatomical confinement was corroborated functionally: photostimulation in the VTA and in the IPN drove opposing behaviors (preference vs. aversion), a double dissociation that would likely be abolished if uncaged agonists diffused appreciably between these regions. The near-UV wavelength required to photolyze the NV cage is strongly attenuated in tissue, limiting the depth and volume a single optical fiber can address. This same property, however, contributes to the spatial confinement documented above: because the excitation light does not travel far from the fiber tip, uncaging remains local. Where deeper or larger-volume release is needed, red-shifted cages absorbing in the visible range or within the tissue optical window would relax the penetration constraint (Ellis-Davies, 2023; Li et al., 2026).

The IPN is activated by aversive experiences (Klenowski et al., 2022; Liang et al., 2024), and optogenetic activation of IPN neurons can drive or enhance avoidance behavior (Wolfman et al., 2018; Liang et al., 2024; Ciscato et al., 2025a). Photorelease of EPI in the IPN evoked place aversion, providing direct evidence that activation of endogenous IPN nAChRs is sufficient to drive avoidance. This aversion was strongly enhanced by a single prior administration of nicotine, providing convergent evidence from an orthogonal approach that the aversive weight of IPN activity is gated by recent nicotine experience (Morton et al., 2018).

Photoactivation in the PAG increased mechanical sensitivity, which is, to our knowledge, the first demonstration of a pronociceptive nicotinic action within the brain. The opposite behavioral effects of systemic and local ABT594 emphasize that the circuit location of receptor activation can determine the sign of a drug’s behavioral effect. ABT594 was developed as an α4β2-preferring nicotinic analgesic and produces antinociception in preclinical models (Decker et al., 1998; Donnelly-Roberts et al., 1998; Holladay et al., 1998; Hayashi et al., 2017). However, systemic administration engages receptor populations across spinal, brainstem and supraspinal sites. The PAG itself is not a uniform analgesic node: distinct PAG neuronal populations can either suppress or facilitate nociception (Samineni et al., 2017), and an antinociceptive vlPAG circuit mediated by α7 nAChRs has been described separately (Sullere et al., 2023). ABT594, however, engages α4β2 rather than α7 receptors, which enhance rather than suppress GABAergic transmission in PAG neurons (Nakamura and Jang, 2010; Sullere et al., 2023). We speculate that focal ABT594 engages α4β2 receptors preferentially on local GABAergic elements, increasing inhibitory tone onto PAG output neurons and suppressing descending antinociceptive drive, the opposite net effect of the α7 nAChR-mediated circuit, despite both acting through nicotinic receptors in the same structure. This is consistent with evidence that direct activation of vlPAG GABAergic neurons produces hypersensitivity (Samineni et al., 2017; Xie et al., 2023). The local pronociceptive effect we observed does not contradict the analgesic profile of systemic ABT594, rather it reveals a local circuit effect that is normally averaged out by concurrent effects in other sites.

Importantly, the method developed here applies to any target for which a potent, caging-tolerant ligand exists. Neither agonist is fully subtype-selective: both bind heteromeric α4β2 and α3β4 nAChRs with high affinity and α7 only weakly (Kesingland et al., 2000; Meyer et al., 2006). Additionally, EPI, unlike ABT594, activates muscle-type nAChRs (Sullivan et al., 1994), making NV-ABT594 the better choice where neuromuscular activation would confound the readout. Caging a subtype-preferring ligand would add selectivity the present probes lack. The same approach extends to antagonists (McClain et al., 2023), where photorelease would locally silence rather than drive endogenous signaling. Extending this approach to other receptors therefore depends mainly on ligand chemistry rather than on biology, making caged high-affinity ligands a broadly applicable strategy for optically targeted neuropharmacology.

More broadly, the ability to restrict the action of a drug to a defined anatomical target reframes what a caged therapeutic could offer. Much of the toxicity of drugs arises not from the target receptor itself but from its activation at unwanted sites; ABT594 is a paradigmatic case, in which genuine analgesic efficacy could not be separated from dose-limiting adverse effects during clinical development (Meyer, 2006). A systemically inert precursor that is converted to an active agonist only where light is delivered offers a route to spatially restricted “smart” pharmacology: the therapeutic action confined to the intended circuit while the rest of the brain and periphery is exposed only to the inactive form (Lerch et al., 2016; Paoletti et al., 2019). Photopharmacology is no longer a purely preclinical proposition: a chemical photoswitch we developed (Mourot et al., 2011) has entered clinical trials, with phase I results reported (Casson et al., 2026) and a phase II trial ongoing. The present work remains a proof of concept in mice; translating a systemically delivered caged agonist to humans would require optimized pharmacokinetics, safe light doses, and a favorable safety profile for the precursor and its photoproducts. Still, as tools for dissecting where in a circuit the beneficial and adverse effects of a drug arise, caged high-affinity agonists offer a route to widening its therapeutic window.

## Methods

### EXPERIMENTAL ANIMALS

Experiments were conducted using adult male and female wild type C57BL/6NRj mice (Janvier labs, France, RRID:IMSR_RJ:C57BL-6JRJ). Mice were housed at 18-23°C under a 12-hour light/dark cycle, with food and water available ad libitum. For behavioral test, all mice were habituated to the experimenter for 1-2 days before the first test day and transported to the testing room at least 1 hour before any procedures to facilitate adaptation to the surroundings. All tests were conducted during the light phase. All procedures for animal maintenance, surgery and behavioral tests used protocols that were in accordance with the recommendations for animal experiments issued by the European Commission directives 2010/63 and approved by Paris Sciences et Lettres (PSL) University, the Hérault department Veterinary Direction, France and the French ministry for higher education, research and innovation.

Animal cohorts: Patch-clamp electrophysiology: n = 5 females, n = 5 males. *In vivo* electrophysiology: n = 8 for NV-EPI, n = 7 males for NV-ABT594. fUSi: n = 3 males. Fiber photometry: n = 44 males. Open-field tests: n = 39 males for NV-EPI; n = 21 males for NV-ABT594. RTPP tests: n = 26 males for the VTA experiment; n = 3 females, n = 22 males for IPN experiments. Von Frey test: NV-ABT594 at 4000 µg/kg n = 13 males; NV-ABT594 at 800 µg/kg n = 8 males; ABT594 at 120 µg/kg n = 6 males.

### CHEMICALS

#### Chemical synthesis and characterization

Commercial reagents were used as received. Epibatidine hydrochloride was obtained from Sigma (E1145-5MG), ABT594 HCl ((R)-5-(2-azetedinylmethoxy)-2-chloropyridine hydrochloride, also known as Tebanicline) was obtained from Tocris (#6576), nicotine hydrogen tartrate salt (nicotine) was obtained from Glentham LifeSciences (GL9693). Synthesis of NV-Epibatidine (**Fig. S2a**): to a solution of epibatidine hydrochloride (10 mg, 0.041 mmol) and nitroveratryl bromide (34 mg, 0.12 mmol) in DMF was added potassium carbonate (17 mg, 0.012 mmol). The mixture was stirred at RT for 48 hours. Water was added to the reaction mixture, which was extracted with ethyl acetate. The organic layer was condensed by under reduced pressure, and purified by flash chromatography to give NV-EPI (14 mg, 0.036 mmol) in 90% yield.

HR-AM: C_20_H_22_ClN_3_O_4_ requires 403.1299, 404.1386, M+1 found.

NV-ABT594 was synthetized as in (Passlick et al., 2018) and stored as powder at −20°C.

#### Experimental solutions

Nicotine hydrogen tartrate salt was dissolved in sterile saline (0.9% NaCl prepared in Milli-Q water). The pH was adjusted to 7.5, and solutions were passed through a 0.2-µm sterile nylon syringe filter (Nalgene, 25 mm). Nicotine aliquots were stored at −20 °C and thawed on the day of use. Nicotine doses are reported as nicotine free-base equivalents.

NV-EPI and NV-ABT594 were handled under red light to minimize unintended photolysis. NV-EPI was dissolved in Milli-Q water to prepare a 500 µM stock solution, aliquoted in 1 ml volumes, and stored at −20 °C. On the day of the experiment, a stock aliquot was thawed and diluted in sterile saline to the required final concentration, sonicated, and protected from light until injection. Fresh working solutions were prepared daily and discarded after use. Stock solutions of NV-ABT594 were prepared at 100 mM in anhydrous DMSO, aliquoted (50 µl), and stored at −80 °C. The stock was diluted in phosphate-buffered saline (PBS) and sonicated before use. Fresh working solutions were prepared daily, protected from light, and discarded after use.

For i.p. injections, compounds were administered at 100 µl per 10 g body weight. Injections were administered using low-dead-volume BD Plastipak syringes (0.5 or 1 ml).

#### Molecular docking

Molecular docking was performed using a high-resolution structure of the α4β2 nicotinic acetylcholine receptor in an agonist-bound conformation (PDB ID: 8ST0). Protein preparation included removal of non-relevant ligands, addition of hydrogens, protonation state assignment at pH 7.4, and energy minimization using the Adopted Basis NR algorithm to a final gradient of 0.001.

Ligand structures for epibatidine, ABT594, NV-EPI, and NV-ABT594 were built and energy-minimized in Discovery Studio 2026. Flexible ligand-rigid protein docking was performed using CDOCKER implemented in Discovery Studio 2026 (Wu et al., 2003). Random ligand conformations were generated from the initial ligand structure through high-temperature molecular dynamics. Docking grids were centered on the orthosteric binding pocket defined by the co-crystallized agonist, with a binding site sphere large enough to accommodate the NV group for the caged derivatives.

### SLICE ELECTROPHYSIOLOGY

#### Slice preparation

Acute brain slices containing the MHb or VTA (320 µm thick) were prepared from C57BL/6J mice aged postnatal day 25-35. Mice were deeply anesthetized with ketamine/xylazine (120 mg/Kg and 10 mg/Kg respectively) and transcardially perfused with an ice-cold sucrose-based cutting solution containing (in mM): 220 sucrose, 11 glucose, 2.5 KCl, 1.25 NaH₂PO₄, 25 NaHCO₃, 7 MgSO₄, and 0.5 CaCl₂. The brain was rapidly removed, and slices were prepared using a vibrating microtome (VT1200S, Leica Microsystems).

Freshly cut slices were transferred to artificial cerebrospinal fluid (ACSF) maintained at 34°C and containing (in mM): 125 NaCl, 2.5 KCl, 2 CaCl₂, 1 MgCl₂, 1.25 NaH₂PO₄, 25 NaHCO₃, and 15 glucose. After an initial recovery period of 15-20 minutes, slices were maintained at room temperature and used for recordings within 6 hours.

### Patch-clamp electrophysiological recordings

Whole-cell patch-clamp recordings were obtained from MHb or VTA neurons at 33-35°C using a MultiClamp 700B amplifier (Molecular Devices). Neurons were visualized using differential interference contrast optics. Recording pipettes were fabricated from thick-walled glass capillaries (1.85 mm outer diameter, 0.84 mm inner diameter; 1B150F-4, World Precision Instruments) and fire-polished to a resistance of 3.5-5 MΩ. For voltage-clamp recordings, pipettes were filled with an intracellular solution containing (in mM): 90 CsMeSO₃, 10 EGTA, 40 HEPES, 4 MgCl₂, 5 QX-314, 2.5 CaCl₂, 10 Na₂-phosphocreatine, 0.3 MgGTP, and 4 Na₂ATP. The solution was adjusted to pH 7.3 with CsOH and to an osmolarity of approximately 300 mOsm. Nicotinic receptor-mediated currents were evoked by local pressure application of nicotine. Where indicated, NV-EPI or nicotinic receptor antagonists were applied through the bath solution. Nicotine (20 µM) was puffed locally at the slice surface near the recorded neuron through a glass pipette (2-3 MΩ) using a pneumatic drug-ejection system (PicoPump PV 820, World Precision Instruments, Germany). Full-field photolysis was achieved with 100-ms pulses from a 365-nm LED (Cairn Research, UK) mounted on the epifluorescence port of the microscope (Olympus); light was directed through the objective (60x) and controlled with an OPTOLED LITE LED driver (Cairn Research, UK) via external voltage modulation. The pneumatic drug-delivery and full-field illumination configurations were adapted from (Bossi et al., 2022). For current-clamp recordings, the intracellular solution contained (in mM): 135 K-gluconate, 5 KCl, 10 HEPES, 0.01 EGTA, 10 Na₂-phosphocreatine, 4 MgATP, and 0.3 NaGTP. The pH was adjusted to 7.3 with KOH and the osmolarity to approximately 295 mOsm. Membrane potential was maintained near −70 mV by injecting a small holding current when necessary. Membrane potentials were not corrected for the estimated liquid-junction potential of approximately 10 mV.

Series resistance was monitored using a −5-mV test pulse during voltage-clamp recordings. Cells were excluded if series resistance exceeded 20 MΩ or changed by more than 20% during the experiment. Current-clamp recordings were excluded if input resistance varied by more than 25% from its initial value. Signals were low-pass filtered at 10 kHz and digitized at 100 kHz using an NI USB-6259 interface (National Instruments, Austin, TX, USA). Data were acquired using Nclamp software (Rothman and Silver, 2018) running in Igor Pro (WaveMetrics, Lake Oswego, OR, USA).

### GENERAL SURGICAL PROCEDURES AND POSTOPERATIVE CARE

The mouse was first placed in an induction chamber and deeply anaesthetized with oxygen (1 L/min) containing 3% isoflurane (Piramal Healthcare, UK) regulated with an anesthesia station (MiniHUB V3, TEMSEGA, France). The top of the skull was cleaned with an iodine solution (betadine gel) and the mouse was then placed in a stereotaxic frame (David Kopf Instruments, CA, USA) with a digital manipulator arm (Stoelting, IL, USA) on a feedback-controlled heating pad maintained at 37°C (RWD ThermoStar System, Shenzhen, China). During surgery, anaesthesia was maintained with 1% isoflurane in oxygen. Ophthalmic ointment was applied to the eyes to prevent drying. The scalp was shaved and a local anesthetic (Lurocaine®, 100 μL at 0.67 mg/kg, s.c.) was applied at the location of the scalp incision, and meloxicam (Métacam, 100 µl at 5 mg/kg, s.c.) was administered before the procedure. At the end of the surgery, an analgesic solution of buprenorphine (Buprecare®, 1 mg/kg) was injected subcutaneously to prepare awakening and supplemented the following recovery days if necessary. The mouse was monitored until it completely recovered and thereafter placed in the homecage.

### GENERAL TISSUE PROCESSING AND IMMUNOSTAINING

The mice were anesthetized deeply with Euthasol (pentobarbital sodium and phenytoin sodium) and transcardially perfused with 1× phosphate-buffered saline (PBS; pH 7.4) followed by 4% paraformaldehyde (PFA) in PBS (0.1 mM; pH 7.4). The perfused brain was removed from the skull and postfixed with 4% PFA in 0.1 mM PBS at 4°C overnight. For vibratome cutting, the brain was thoroughly washed in 1× PBS and thereafter sectioned coronally (50-μm thickness) using a vibratome (Leica VT1000S, Leica Microsystems, Nussloch GmbH). The sections were stored in 1× PBS at 4°C. Vibratome cut (free-floating) sections were permeabilized for 1 hour in a blocking solution of 1× PBS containing 3% bovine serum albumin (BSA, Sigma; A4503) and 0.2% Triton X-100 at room temperature (RT), and thereafter incubated with primary antibodies (see antibody list) in 1× PBS containing 1.5% BSA and 0.2% Triton X-100 at RT for 12-24 h. Then the sections were rinsed three times in 1× PBS and incubated with a species-specific fluorophore-conjugated secondary antibody in a solution of 1.5% BSA in PBS for 3-5 hours (see antibody list for details). Slices belonging to mice with optical fibers implants in the IPN were additionally incubated with a Cy3 anti-mouse secondary antibody to help visualize the fiber tract. After three rinses in PBS (10 minutes each), slices were mounted on glass slides (Superfrost Plus, ThermoFisher Scientific), and mounted using Prolong Gold Antifade Reagent with DAPI (Invitrogen, P36931). Images were acquired at 5, 10 or 20× magnification with an epifluorescent microscope (Zeiss Axio Imager M1), and images captured using a camera (Zeiss AxioCam MRm) and preprocessed with Fiji (RRID:SCR_002285) (Schindelin et al., 2012).

### Primary antibodies used

Chicken anti-GFP antibody (Sigma-Aldrich, AB16901; RRID:AB_90890) at 1:1000 dilution. Sheep polyclonal anti-Tyrosine Hydroxylase (Sigma-Aldrich, AB1542; RRID:AB_90755) at 1:500 dilution.

### Secondary antibodies used

Alexa488-conjugated anti-chicken secondary antibodies (Invitrogen, A-11039; RRID:AB_2534096) at 1:1000 dilution. Cy3 anti-mouse secondary antibody (Invitrogen, M30010; RRID:AB_11181146) at 1:500 dilution. Alexa Fluor 647-conjugated anti-Sheep secondary antibodies (Jackson ImmunoResearch, 713-605-003; RRID:AB_2340750) at 1:1000 dilution.

### ACUTE *IN VIVO* EXTRACELLULAR ELECTROPHYSIOLOGY

#### Acute preparation, venous catheterization and craniotomy

Mice were prepared as in General Surgery procedure. For experiments involving i.v. administration of saline, nicotine or caged compounds, the saphenous vein was catheterized using a 30G needle connected to polyethylene tubing (PE10) and a Hamilton syringe (Ciscato et al., 2025b). The scalp was opened and a craniotomy was drilled above the target structure. A second craniotomy was made above the cerebellum to insert a ground/reference screw.

Extracellular activity was recorded from either the ventral tegmental area (VTA) or the interpeduncular nucleus (IPN). VTA recordings were targeted using stereotaxic coordinates approximately 3.0-3.4 mm posterior to bregma, 0.3-0.6 mm lateral to the midline, and 4.0-4.8 mm below the brain surface. IPN recordings were targeted with a 5° angle using coordinates approximately 3.3-3.6 mm posterior to bregma, 0.35-0.5 mm lateral to the midline, and 4.2-4.9 mm below the brain surface. Final coordinates, probe angle and probe identity were recorded for each session and stored in the experiment metadata table.

Electrophysiological signals were acquired using a 32-channel silicon neural probe (ASSY-37 H10b, Cambridge Neurotech, UK) coupled to an optical fiber (100 μm core, high OH, 0.39NA, 0.7mm taper-tip Lambda-b fiber; Optogenix, Italy). This optrode was connected via an acute probe adapter (32 channels Samtec to Omnetics, ADPT A32-Om32, Cambridge Neurotech) to an HS-36-PTB Omnetics headstage (Neuralynx). The headstage was connected through a shielded tether (TETH-HS-36-Litz-3M, Neuralynx) and an ADPT-HS-36-DRS adapter to a Digital Lynx 4SX acquisition system (Neuralynx). Signals were digitized and recorded using Cheetah 5.0 software (Neuralynx) and monitored online via NeuroExplorer (v5.430; Nex Technologies). The electrode reference was connected to the cerebellar ground screw. After lowering the probe into the target structure, the probe was allowed to stabilize for approximately 30 minutes before recordings began. Signals were sampled at 32 kHz and saved as spike-band data after hardware/software band-pass filtering between 600 and 6000 Hz. In this dataset, only spike-band signals were available for analysis; local field potential signals were not used. Spike waveforms appeared as positive-going deflections in the recorded data.

Optical stimulation was delivered using a 365-nm LED source (Optogenetics-LED-365, Prizmatix) triggered by a data acquisition board (BNC-2110, National Instruments) controlled by a custom LabVIEW program (National Instruments). Light stimulation consisted of pulses of variable duration, from 1 s to 30 s epochs, delivered at 8 mW. Optical power was measured at the fiber tip before each experiment using a 1916-R power meter with an 818-ST2-UV/DB UV-enhanced silicon detector (200 - 1100 nm; Newport).

After a 3-minute baseline recording, mice received i.v. saline injection followed by control light stimulations (minimum three). Recordings continued for 5 minutes after saline administration. Then, NV-EPI was administered either i.v. (50 µg/kg) or i.p. (100 µg/kg). Intravenous injections of saline and NV-EPI were delivered at 10 µl per 25 g body weight, whereas i.p. saline or NV-EPI injections were delivered at 100 µl per 10 g body weight.

#### Silicon probe track reconstruction

To reconstruct probe location, the silicon probe was coated with CM-DiI (1,1′-dioctadecyl-3,3,3′3′-tetramethylindocarbocyanine perchlorate; Invitrogen V22889), a fixable lipophilic dye. The probe shank was painted repeatedly with a small drop of CM-DiI and allowed to dry until visibly coated. After recording, brain sections containing the fluorescent probe track were imaged using a Zeiss epifluorescence microscope at 5× magnification. The sections were registered to the Allen Common Coordinate Framework version 3 (Allen CCFv3), and probe trajectories were reconstructed using the DMC-BrainMap pipeline (Jung et al., 2026). The anatomical location of each unit was assigned according to the position of the probe contact with the largest absolute template amplitude for that unit.

#### Spike sorting and manual curation

Spike sorting was performed with Lupin (Garcia et al., 2026) (v2026.01), an internal template-matching sorter in SpikeInterface (Buccino et al., 2020) (v0.104.x). The 32 Neuralynx spike-band channels were concatenated into an interleaved int16 binary file and loaded with the Cambridge Neurotech H10b probe geometry. The session-specific Neuralynx ADBitVolts value was read from metadata and used to scale signals to microvolts. Since spikes were positive-going in our recording configuration, signals were inverted before sorting. No additional filtering was applied because recordings were already band-pass filtered between 600 and 6000 Hz at acquisition. A global median common average reference was applied before inversion. Lupin was run on the full continuous recording using negative peak detection, a threshold of 5 median absolute deviations, no motion correction, and a waveform window of 0.5 ms before to 1.5 ms after the spike. After Lupin sorting, we used the SpikeInterface SortingAnalyzer to compute unit templates, waveforms, spike amplitudes, noise levels and principal components. Units were inspected interactively using the SpikeInterface GUI (sigui). First, to facilitate manual curation, units were first automatically pre-labelled using region-aware quality-control rules. Units with fewer than 50 spikes, peak template amplitude below 20 µV, or ISI violation rate ≥ 15% for ISIs < 2 ms were pre-labelled as noise. Units with ISI violation rates between 2% and 15% were pre-labelled as multi-unit activity. Units with ISI violation rates < 2%, at least 100 spikes and peak template amplitude ≥30 µV were pre-labeled as good candidates. These automatic labels were used only to prioritize manual review and were not used as final acceptance criteria. Because IPN neurons may show high tonic firing rates, firing rate was not used as an exclusion criterion; high firing rates were instead flagged for manual inspection. For final unit classification, we used a custom Jupyter-based curation workflow, which displayed for each unit a panel with waveform overlays, ISI histogram, autocorrelogram, spatial footprint across the H10b contacts, and firing rate over the session with event markers. Based on these views and the pre-computed quality metrics (ISI violations, spike count, amplitude), units were manually labelled as good (well-isolated single unit), multi-unit activity (MUA), or noise. Units were classified as good when they showed a stereotyped waveform, a localized spatial footprint on adjacent probe contacts, a clear refractory trough in the autocorrelogram, low ISI violations and stable firing outside expected experimental manipulations. Units with recognizable spike waveforms but incomplete refractory periods or variable waveform features were classified as multi-unit activity. Units with non-physiological waveforms, near-zero amplitudes, global artifacts, light/injection artifacts or unstable templates were classified as noise. Only units labelled as good were used for single-unit analyses. MUA was included only in explicitly indicated population-level analyses. For each curated unit, spike times were exported in seconds relative to recording onset and used for all downstream analyses.

#### Electrophysiology response classification and statistical analysis

Light-evoked responses were classified at the single-unit level by comparing firing during light delivery (0 to T, where T was the stimulus duration) with a duration-matched baseline (−T to 0 s). Units were classified as activated or inhibited when the mean firing-rate change exceeded prespecified thresholds and its direction was consistent across trials. Directional consistency required both trials for units with two trials or at least two-thirds of trials for units with three or more. The effect-size threshold was 0.50 spikes/s and a ≥ 30% change for units with baseline firing ≥ 0.20 spikes/s, or an absolute change of ≥ 0.20 spikes/s for units with lower baseline firing. All other units were classified as unchanged. In drug-plus-light sessions, units not meeting these criteria during illumination were also evaluated from light onset to 60 s after light offset to identify delayed or persistent responses. Light-only controls were classified using the illumination period only. For units tested at multiple durations, one canonical duration was selected according to a predefined duration ranking and trial availability. For NV-EPI experiments, compatibility of i.v. and i.p. administration was assessed separately in the VTA and IPN using linear mixed-effects models of the light-evoked firing-rate change, with mouse as a random intercept and time from injection to illumination included as a covariate (Supplementary Fig. S3b). As no route-associated difference was detected, IV and IP recordings were pooled for subsequent NV-EPI analyses. Baseline-to-light firing-rate changes and differences among response classes were tested using linear mixed-effects models, with mouse as a random intercept and recording session and/or unit included as variance components where appropriate. NV-EPI and NV-ABT594 cohorts were analyzed separately. To assess the specificity of the light response, light-only-control units were pooled across cohorts and deduplicated at the unit-recording level. Equivalence to a negligible effect was tested using two one-sided tests with bounds of ± 0.5 spikes/s and was concluded when the 90% confidence interval of the mean baseline-to-light change fell entirely within these bounds.

### FUNCTIONAL ULTRASOUND IMAGING

#### Cranial-window and VTA-fiber implantation for fUSi

Following the general surgery procedure, a parasagittal scalp incision was made to expose the skull. A 2 mm × 4 mm cranial window was delineated over the imaging region, and the overlying skull was thinned carefully and removed with a dental drill while preserving the dura mater. A 125-µm-thick TPX membrane (Goodfellow) was cut to match the dimensions of the cranial window, disinfected with alcohol, air-dried, positioned over the exposed cortex, and sealed to the skull using dental cement (Superbond Universal Kit, Sun Medical, Japan; GC Unifast TRAD). An 8-mm-long, 200-µm-core optical fiber was mounted on a movable nanodrive (Cambridge Neurotech) and implanted to target the ventral tegmental area (VTA) at a 40° posterior angle (AP, −6.4 mm; ML, ±0.5 mm; DV, −5.5 mm, relative to bregma). A custom permanent fUS baseplate holder was positioned above the imaging plane and centered on the cranial window (13 mm × 3 mm × 5 mm, 0.3 g) and movable silicon protective cap (0.25 g) was positioned to protect the cranial window and implant between recording sessions. The implant was secured using dental cement (GC Unifast TRAD), and diluted povidone–iodine solution (1:10 Betadine) was applied around the incision site. At the end of the surgery, mice received intraperitoneal 5% glucose for fluid replacement and 0.2 ml Metacam (5 mg/kg, s.c.). Buprenorphine (Buprecare, 0.1 mg/kg) and Metacam (Meloxicam, 20 mg/kg) were injected subcutaneously every 12 hours and 24 hours, respectively, during 48 hours. All mice were left to recover for 7 days.

#### fUSi acquisition

fUSi was performed by ultrafast compound Doppler imaging to measure changes in CBV. Images were acquired using a 256-channel Iconeus One ultrasound scanner (Iconeus, France) and a 64-element linear-array ultrasound probe (Iconeus, France) with a 15-MHz central frequency, 0.11-mm element pitch and 46% bandwidth at −6 dB (11-18 MHz). The in-plane spatial resolution was 0.11 × 0.11 mm. The acoustic lens provided an elevation focal distance of 8 mm and an elevation focal width of approximately 400 µm, defining the thickness of the imaging plane. The available field of view was 20 mm x 6.4 mm. fUS images were cropped along the z-axis to remove non-brain voxels. For each compounded frame, the imaging field was insonified using 10 plane waves spanning angles from −5° to +5° in 2° increments. Plane-wave transmissions were delivered at a pulse-repetition frequency of 500 Hz. Power-Doppler images were acquired at 2 Hz, with a 400-ms acquisition period followed by a 100-ms pause. This acquisition scheme permitted prolonged recording without frame loss or memory saturation. A 150-cm cable length and 12-g probe weight ensured relative freedom and comfort for the animal. Mice were recorded in their homecage in freely-behaving conditions. The fUS tethers are attached to a counterweight system that reduces the apparent weight of the ultrasound probe on the animal’s head.

#### Photostimulation sessions

Each recording session lasted 40 minutes during which fUSi data was continuously acquired. Each session included (1) a 5-minute baseline epoch where fUSi data was acquired at rest (2) a 5-minute pre-drug epoch, starting with an i.p. injection of saline (100 µl per 10 g body weight) followed by three 15-s light-stimulation trials, with 1 minute of inter-stimulation interval (3) a 30-minute test epoch, starting of an i.p. injection of NV-EPI (200 µg/kg at 100 µl per 10 g body weight), followed by fUSi acquisition during photorelease. This 30-minute test epoch was composed of a 5-minute pre-stimulation block, followed by received three trials triplet, with an inter-trial interval of 120 s and inter-triplet interval of 150 s, and a final 5-minute baseline recovery block. Triplet differed only by stimulation duration (5, 10 and 15 s, respectively), presented in a randomized order across animals. Optical stimulation of the VTA was delivered using a fiber-coupled 365-nm LED (Optogenetics-LED-365; Prizmatix). Light was transmitted through a 1-m patch cord with a 1,000-µm core and numerical aperture of 0.63 (OG-Fiber-1000; Prizmatix) to a low-friction rotary joint (Prizmatix), and then through a 500-µm-core patch cord (OG-Fiber-500-1.25; Prizmatix) connected to the implanted optical fiber. Optical power was adjusted to 8 mW at the tip of the implanted fiber and was measured before each recording session using a 1916-R power meter equipped with an 818-ST2-UV/DB UV-enhanced silicon detector (Newport). Light pulses were programmed and triggered using a USB pulse-train generator (Pulser Plus; Prizmatix) interfaced with the acquisition computer. Timestamps of video frames and light stimulation pulses were collected and digitized on a data acquisition board (Intan Technologies, USA) and used to synchronize multimodal data a posteriori.

#### fUSi preprocessing and quantification

Power-Doppler images were reconstructed from raw backscattered ultrasound signals. Radiofrequency data were beamformed to generate complex in-phase/quadrature (IQ) data. Tissue-motion and pulsatility-related clutter were removed using singular-value decomposition filtering, in which the first 60 singular components were excluded. Power-Doppler images were then computed from the filtered Doppler signal. This measure reflects the concentration of moving echogenic scatterers within each voxel and was used as an index of relative cerebral blood volume. No motion correction method was applied, as the custom fUS baseplate enables a tight mechanical contact between the probe and the skull, resulting in high-quality low-noise fUS data, even during movement. Functional ultrasound images were manually registered to coronal sections of the Allen Mouse Brain Atlas. The custom fUS baseplate allowed for single-plane imaging at a fixed position, defined at the surgery. The coordinate of the imaging plane along the dorso-ventral axis (Bregma −3.5 mm) was cross-validated by atlas registration using the Paxinos Atlas. The fiber tip location was identified a posteriori during histology and registered onto the fUS imaging plane.

For each voxel, CBV signals were normalized to the 5-minute pre-injection baseline according to the following equation:

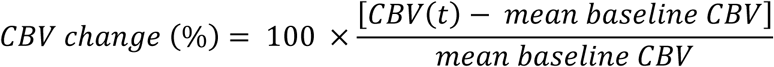

Regional CBV time series were obtained by averaging normalized voxel-wise signals within each region of interest. Regional and whole-brain CBV time courses were smoothed using a Gaussian kernel with a 10-s window width. The resulting datasets consisted of Power-Doppler image sequences acquired at 2 Hz over recording sessions of 40 minutes.

#### fUSi statistical analysis

To compute peri-event regional responses (**Fig. 3d-e**), each regional CBV time series was interpolated and re-aligned to a peri-stimulation time vector starting 10 s before stimulation onset and ending 30 s after, with a timestep of 1 s. Each individual trial response was then normalized by subtracting the mean value over the 10-s pre-stim window, for that trial. Inter-trial mean responses were then computed for all sessions using the first 6 trials, for the test stimulations, so as to include at least one triplet of long (10 or 15 s-long) photostimulations and the first two trials for the control stimulations, as the third and last control trial was too close to the subsequent injection (and thus contained artefacts). The area under curve (AUC) is then computed by summing all positive values and subtracting all negative values of the mean peri-response CBV series, over the −10 s to 30 s peri-stimulation time vector. Our dataset consisted of n = 3 animals including two sessions per animal. We statistically tested for the null hypothesis that peri-stimulus AUC would be drawn from the same underlying distribution in the contralateral and stimulated VTA (and stimulated VTA versus light-control) using a paired two-sided Student t-test and rejected it, using a type I alpha error of 0.5. Statistical robustness was further assessed by nonparametric bootstrap resampling of the recording-level paired differences. For each comparison, paired differences were resampled with replacement 5,000 times, and bootstrap 95% confidence intervals were calculated from the resulting distribution of mean paired differences.

#### Histological/implant verification

Brains from fUS cohorts were collected as described under General tissue processing and immunostaining, sectioned coronally at 50 µm, and all consecutive sections spanning the VTA and adjacent regions were collected, immuno-stained for TH^+^ and mounted. After imaging with a Zeiss epifluorescent microscope at 5x, and the coronal sections were analyzed with FIJI, and optical fibers’ trajectories were reconstructed from the tissue lesion tract, to assign the anatomical location of the tip of the fiber to a specific region. Mice with misplaced fibers were excluded from all analyses.

### FIBER PHOTOMETRY

#### Viral delivery and NAc/VTA fiber implantation

GRAB_DA2m_ (G protein-coupled receptor [GPCR]-activation-based), full name AAV1-hSyn-GRAB_DA2m_ (titrating at 1.23×10^14^ particles/ml), was produced as previously described (Nguyen et al., 2021) by Deniz Dalkara (Institut de la Vision, Paris, France) from plasmid psAAV-hSyn-GRAB_DA2m_ generously gifted by Yulong Li (Sun et al., 2020). For injection of GRAB_DA2m_, a biosafety cabinet class 2 with sterile surgical environment was used. A median incision revealed the skull which was drilled at the level of the targeted structure. Craniotomy was performed over the targeted brain area. Sterile saline 0.9 % was applied regularly over the skull while drilling to remove bone dust and control heat generation. The total diameter of the burr hole was not larger than the diameter of the drill bit (∼100 μm). Fixed to a One-axis Oil Hydraulic Micromanipulator (MO-10, Narishige, Tokyo, Japan), the micropipette (graduated borosilicate glass capillary; Wiretrol I, Drummond, PA, USA) was filled with mineral oil and the virus. The tip of the micropipette was lowered through the dura to reach the target location; 0.5 μl virus was injected at a rate of 0.1 μl/min. The micropipette was kept in place for 5 minutes after each finished injection and thereafter slowly retracted. The skin was closed with Ethilon 5/0 sutures (Ethicon, NJ, USA).

Adult male and female mice were injected with 0.5 μl of AAV1-hSyn-GRAB_DA2m_ in the NAc [anteroposterior (AP): 1.45 mm, mediolateral (ML): ±1.75 mm, dorsoventral (DV): −4.1 mm relative to bregma]. Injected mice were then implanted with optical fibers (200 µm diameter, NA 0.39, Thorlabs) aimed directly above the NAc (same AP and ML coordinates, but DV: −4.0; steel ferrule) and the VTA (AP: 3.10 mm, ML: ±0.5 mm, DV: −4.2 mm relative to bregma; ceramic ferrule). The optical fibers were secured onto the skull using dental adhesive and cement (Superbond Universal Kit, Sun Medical, Japan) and light-curing cement (SpeedCEM Plus, Ivoclar Vivadent).

#### Fiber photometry hardware and signal acquisition

Fluorescence was acquired using a single-site fiber photometry system (Doric Lenses) configured for dual-wavelength operation. For systemic nicotine experiments, the two connectorized LEDs (405 nm and 465 nm; Doric Lenses) were modulated sinusoidally at distinct carrier frequencies (333.8 Hz and 220.5 Hz, respectively) and coupled into a fluorescence mini-cube via fiber patch cords. For caged-ligand experiments, the 405-nm excitation LED was disabled throughout acquisition to minimize inadvertent photolysis because the nitroveratyl (NV) caging group used in this study absorbs minimally at wavelengths up to ∼400-420 nm (photolysis peak ∼350-380 nm).

Excitation light was delivered to the implant through a low-autofluorescence optical fiber interfaced to the implanted cannula via a zirconia sleeve. Emitted fluorescence was collected through the same fiber, routed to a visible-range photoreceiver (AC-coupled, low-gain mode; New Focus Model 2151, MKS Instruments), and digitized via the Doric fiber photometry console. Signals were acquired in Doric Neuroscience Studio (v5.2.2.5 and v6.6.3.0) at approximately 1.2 kHz following lock-in demodulation, with a hardware low-pass filter set to 12 Hz.

#### Photorelease of caged ligands in the VTA

Photolysis of caged compounds in the VTA was achieved with a fiber-coupled 365-nm ultra bright LED (Optogenetics-LED-365; Prizmatix). Light was transmitted from the LED source through a 1,000 µm core, NA = 0.63, 1 m patch cord (OG-Fiber-1000; Prizmatix) to a low-friction rotary joint (Prizmatix), and from the rotary joint to the implanted fiber cannula via a 500 µm core patch cord (OG-Fiber-500-1.25; Prizmatix), permitting unrestricted animal movement. Stimulus trains were programmed and triggered using a USB pulse-train generator (Pulser Plus; Prizmatix) interfaced with the acquisition computer.

#### Preprocessing and event-aligned analysis

Photometry data were exported as Doric CSV or HDF5 (.doric) files and processed in R using custom scripts. Raw demodulated 465 nm channels were down-sampled by a factor of 100 after discarding the initial 100 samples, yielding an effective sampling rate of ∼12 Hz. The down-sampled trace was denoized with a zero-phase 4th-order Butterworth low-pass filter (1 Hz cut-off; forward–reverse filtfilt implementation), which suppresses high-frequency noise without introducing temporal distortion. To correct for slow photobleaching and baseline drift, a constrained double-exponential function was fitted to the smoothed 465 nm signal and subtracted; when no satisfactory exponential fit was obtained, a linear detrend was applied as a fallback. The resulting bleach-corrected 465 nm signal was used for all ΔF/F and z-score analyses.

For NV-EPI duration and power experiments and for 10-15 s NV-ABT594 stimulations, a local pre-stimulus baseline window (typically −5 to 0 s) was defined for each TTL-detected stimulation epoch, and responses were expressed as ΔF/F or z-scores relative to this baseline. Stimulation events were detected from the digital TTL channel, and isolated events of the desired duration (e.g. 1, 5, 10, 15 s) were aligned to time 0. For each event we computed the z-score time-course, the peak z-score, and the area under the z-score curve (AUC) over a stimulation-specific analysis window (e.g. 0-2 s for 1 s pulses, 0-20 s for 5-15 s stimulations). Per-mouse means and group-level statistics were then derived from these event-level metrics using linear mixed-effects models and Welch t-tests as appropriate.

For short NV-ABT594 stimulations (0.25-1 s), where heavy filtering and global detrending can attenuate fast transients, a supplementary raw-based analysis was performed. Raw 465 nm signals and raw TTL traces were used to identify 0.5-2.0 s TTL pulses, and event-aligned z-score traces were computed using only a local baseline (−5 to 0 s before each stimulus) without global bleach correction. AUC and peak z-scores were quantified over a 0-2 s window and compared between NV-ABT594-injected and light alone control mice, as reported in the main text.

#### Systemic nicotine photometry preprocessing

For systemic nicotine (0.5 mg/kg i.p.) experiments, we acquired dual-wavelength, fiber-photometry signals (465 nm and 405 nm) from GRAB_DA2m_-expressing NAc fibers using the same Doric console. Raw demodulated traces were first down-sampled by a factor of 10 to ∼120 Hz and cropped to remove initial artefacts. To remove instrument autofluorescence, we subtracted the mean 465 nm and 405 nm values recorded in a separate autofluorescence session from the corresponding channels. Motion and hemodynamic artefacts were then corrected by fitting the 405 nm channel to the 465 nm channel with a simple linear regression and subtracting the fitted 405-scaled component from the 465 nm trace, followed by adding back the original 465 nm mean offset. For recordings with residual slow photobleaching, we optionally fitted a single- or double-exponential function to the 465 nm baseline and subtracted the fitted trend prior to 405-based motion correction. After motion/bleach correction, we extracted a long window around each i.p. nicotine injection (typically −300 to +1200 s relative to injection time) and computed ΔF/F and z-score traces using the mean and standard deviation of the 300 s pre-injection baseline as F₀ and σ₀. For each mouse we then quantified the peak z-score and AUC over 0-600 s, as well as onset latency, time-to-peak, and half-decay time for comparison with the NV-EPI and NV-ABT594 uncaging responses. One mouse was excluded from the analysis due to a fiber-cabling issue.

#### Fiber Photometry data analysis

All analyses were performed in R (v4.6.0) via RStudio (v2026.04.0+526; Posit PBC) using custom scripts (deposited at 10.5281/zenodo.23071073) and standard packages (lme4, lmerTest, emmeans, ggplot2). Stimulation events were detected from the digital TTL channel using a threshold-based rising-edge detector, and each pulse was characterized by its onset time, duration, and inter-pulse intervals. Isolated events were defined as those with no other TTL pulse within a user-defined gap (typically ≥20 s for duration ladders and ≥10 s for repeatability protocols), and all analyses were restricted to isolated pulses at the desired durations (e.g. 1, 5, 10, 15 s for NV-EPI; 0.25-10 s for NV-ABT594). For each isolated pulse, an event-aligned segment of the bleach-corrected 465 nm signal was extracted using a pre-stimulus window (t = −5 to 0 s) and a post-stimulus window tailored to the expected response kinetics (e.g. t = 0-20 s for 10-15 s stimulations, t = 0-10 s for shorter pulses). A local baseline mean and standard deviation were computed from the pre-stimulus window, and responses were expressed as z-scores relative to this baseline. Three summary metrics were quantified per event: (i) the full z-score time-course, (ii) the peak z-score within a predefined analysis window, and (iii) the area under the z-score curve (AUC) over that window (e.g. 0-2 s for 1 s pulses, 0-10 s for intermediate durations, 0-20 s for 10-15 s pulses). For NV-EPI duration and power experiments, event-aligned z-score traces were averaged across events within each mouse to obtain per-mouse mean traces, from which per-mouse AUC and peak values were derived for each duration or power level. Group-level summaries are reported as mean ± SEM across mice. Statistical comparisons across durations or power levels were performed using linear mixed-effects models of the form:

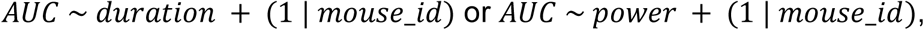

with identical models for peak, treating the fixed effect as a categorical factor for omnibus tests or as a numeric predictor for trend analysis. Model fits were obtained using lme4 and lmerTest with Satterthwaite degrees of freedom; significance of fixed effects was assessed by Type III ANOVA. Where appropriate, post-hoc comparisons (Dunnett contrasts versus the shortest duration or lowest power level) were performed, and key results were verified using Welch’s t-tests on per-mouse means.

For NV-EPI repeatability experiments (0-7 hours and 24 hours), single-duration (15 s) stimulations were analyzed across time points. Per-mouse AUC and peak were computed at each time point, and time-dependence was assessed with linear mixed-effects models of the form:

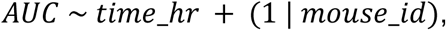

where time_hr denotes hours since injection, with analogous models for peak. Per-time point one-sample t-tests (AUC > 0 and peak > 0) were used to identify time points at which uncaging remained effective, and Dunnett’s post-hoc tests were used to compare each time point against time 0 h.

For NV-ABT594 duration ladders (0.25-10 s), the same event-detection and z-score pipeline was applied with shorter analysis windows (e.g. 0-3 s for 0.25-1 s pulses, 0-10 s for longer pulses) to account for the faster kinetics of NV-ABT594-evoked dopamine release. Duration dependence was assessed using linear mixed-effects models analogous to those for NV-EPI. For NV-ABT594 repeatability protocols (0-5 hours post-injection), series of 1 s stimulations were analyzed with a 0-2 s AUC window; mixed-effects models of AUC and peak versus time_hr were fitted, followed by per-time point one-sample t-tests and Dunnett contrasts versus time 0 hour to identify when response amplitude diverged significantly from the initial value. For responses to very short pulses, a raw-based analysis was additionally performed as described above.

For all comparisons, normality of per-mouse means was assessed by Shapiro–Wilk tests; where necessary, key results were confirmed with non-parametric tests (Wilcoxon rank-sum or signed-rank). Unless otherwise indicated, data are reported as mean ± SEM and two-sided *p* values < 0.05 were considered statistically significant.

To quantify decay kinetics, we fitted a mono-exponential function to the early decay phase using nonlinear least squares (nls, R). For each trial, we then normalized the z-scored trace to its own maximum over the 0-120 s post-stimulus period and averaged normalized traces first within mouse and then across mice to obtain group-mean responses. For NV-EPI 15 s, the fit was restricted to the post-stimulus peak and the following 20 s; for NV-ABT594 1 s, the fit was restricted to the early peak and the following 50 s. The fitted time constant (τ_decay) from each trial was averaged per mouse and compared between NV-EPI and NV-ABT594 using event-level linear mixed-effects models with stimulation mode as fixed effect and mouse as random intercept.

For comparison with systemic nicotine, one response per mouse was analyzed for NV-EPI (15 s light stimulation), NV-ABT594 (1 s light stimulation), and nicotine (0.5 mg/kg, i.p.). Responses were z-scored relative to local pre-stimulation baselines for caged-ligand experiments and to the 300 s pre-injection baseline for nicotine. Peak amplitude, time to peak, and post-stimulation half-decay time were quantified; half-decay time was defined as the interval from the local post-stimulation peak to the first sustained decline below 50% of that peak. Per-mouse metrics were compared using mixed-effects models with mode as a fixed effect and mouse identity as a random intercept; singular models were analyzed using one-way linear models with Tukey-adjusted post-hoc comparisons.

#### Histology and verification of fiber placement and viral expression

Brains from fiber photometry cohorts were collected as described under General tissue processing and immunostaining, sectioned coronally at 50 µm, and all consecutive sections spanning the NAc and VTA and adjacent regions were collected, immuno-stained for green fluorescent protein (GFP) or tyrosine hydroxylase (TH^+^) and mounted. After imaging with a Zeiss epifluorescent microscope at 5x, the coronal sections and the coronal sections were analyzed with FIJI, and optical fibers’ trajectories were reconstructed from the tissue lesion tract, to assign the anatomical location of the tip of the fiber to a specific region. Viral expression level and specificity were evaluated manually; animals with misplaced fibers or absent or non-specific viral expression were excluded from all analyses.

### BEHAVIOR

#### Open-field locomotor assay

To assess the degree to which caging protects against the locomotor side effects of nicotinic agonists, we performed open-field locomotor assays using a circular open-field arena (40 cm diameter) illuminated at 100 lux. Mice received i.p. injections of EPI (0.2, 2, or 10 µg/kg), NV-EPI (0.2, 2, 20, or 200 µg/kg), ABT594 (4 or 40 µg/kg), NV-ABT594 (0.8, 8, 80, 800, or 4000 µg/kg), or saline (vehicle control) at a volume of 100 µl/10 g body weight. Five minutes after injection, mice were placed in the open field and allowed to explore freely for 30 minutes.

To assess the reversibility of the transient locomotor side effect observed at the highest NV-EPI dose, a separate cohort of mice (n = 7 males) received NV-EPI (200 µg/kg, i.p.) and were placed in the open field 1 hour after injection for 30 minutes, and their locomotion was compared with saline-injected controls (n = 7 males) and with animals tested 5 minutes after injection (n = 7 males).

Behavioral data were collected using an overhead video camera recording at 30 frames per second, connected to a custom video-tracking system (code available at 10.5281/zenodo.23071073). Animal position (x, y coordinates) and frame-by-frame displacement were extracted using EzTrack (RRID:SCR_021496) (Pennington et al., 2019). Total distance traveled (cm), mean distance per 1-minute bin, and speed (cm/s) were computed from the EzTrack output using custom Python scripts (code available at 10.5281/zenodo.23071073).

Representative locomotor traces were selected as the animal within each group whose total distance was closest to the group mean. Dose-response relationships were modeled using a constrained four-parameter logistic function fitted to total distance traveled during the 30-minute open-field session. The Hill slope was fixed at 1; therefore, the model estimated three parameters: the upper asymptote, lower asymptote and EC_50_. The model was defined as:

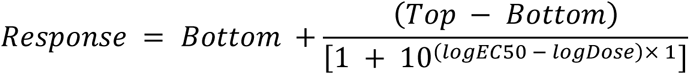

To constrain the fitted curve at the limits of the dose range, two artificial anchor points were included: a nominal low dose of 1 × 10⁻⁵ µg/kg, assigned the maximum observed locomotor response corresponding to the saline-treated animal, and a nominal high dose of 2 × 10⁶ µg/kg, assigned a response of zero. These anchor points represent the assumptions that locomotor activity approaches its maximum in the absence of drug and approaches zero at extremely high doses. EC_50_ values were estimated from the fitted models using the ED() function from drc Package. To examine the reversibility of the acute locomotor effect of NV-EPI, a separate cohort of male mice received NV-EPI (200 µg/kg, i.p.) and was tested in the open field 1 hour after injection. Total distance traveled during the 30-minute session was compared with that of saline-treated controls and NV-EPI (200 µg/kg, i.p.) injected mice tested after 5 minutes using a linear model with treatment group as a fixed effect. Pairwise comparisons were estimated using marginal means and adjusted for multiple comparisons using the Bonferroni method.

#### Real-time place preference (RTPP) and aversion assays

For the VTA experiments, an optical fiber was implanted targeting the VTA (coordinates: AP = −3.1 mm, ML = ± 0.5 mm, DV = −4.2 mm, all from Bregma). For the IPN experiments, mice carried an optic fiber targeting the IPN (coordinates: AP = −3.4 mm, ML = ± 0.4 mm, DV = −4.6 mm, all from Bregma, angle 5°). The optical fibers were secured onto the skull using dental adhesive and cement (Superbond Universal Kit, Sun Medical, Japan) and light-curing cement (SpeedCEM Plus, Ivoclar Vivadent).

RTPP test was performed in a room illuminated with red shifted lamps. It consisted of a custom-build white plastic arena (60 × 30 × 30 cm; ESPCI Paris workshop), comprised by two indistinguishable chambers. The test went on for 15 min. One chamber was paired with light stimulation of 200 ms pulses at 2 Hz (200 ms ON/300 ms OFF) during the entire period that the animal stayed in the stimulus-paired side. The choice of paired chamber was counterbalanced across mice. Animals were placed in the no-stimulation chamber at the start of the session and light stimulation started at every entry into the paired chamber. LED (365 nm; Prizmatix) stimulation was controlled by Bonsai software (Lopes and Monteiro, 2021) (workflow available at 10.5281/zenodo.23071073), interfaced with PulserPlus and an Arduino Uno R3 microcontroller (RRID:SCR_017284) for precise patterned stimulation. LED patterns were predetermined using PulserPlus software. Animal activity was recorded using a video camera connected to Bonsai, recording trajectories at 30 fps.

For standard VTA and IPN RTPP experiments, mice received NV-EPI (200 µg/kg, i.p.) 120 minutes before the behavioral session. For nicotine-primed RTPP/RTPA experiments, mice received nicotine (0.5 mg/kg, i.p.) and NV-EPI (200 µg/kg, i.p.) 120 minutes before testing, as in (Morton et al., 2018). All i.p. injections were administered at 100 µl per 10 g body weight.

Time spent in each chamber, as well as total distance traveled (m), speed, fraction of time spent moving, entry counts, latency to first entry into the stimulated chamber, and bout duration, were measured by ezTrack and custom-scripts in R. Occupancy of the stimulation-paired chamber was quantified as % time in stimulated side and as derived preference metrics. Preference (%) was defined as the difference between times spent in the stimulated and non-stimulated chambers, normalized to session duration. For VTA RTPP, the principal endpoint shown in the main figure was preference across the full 15-minute session, whereas for nicotine-primed IPN RTPA the principal endpoint shown in the main figure was preference during the first 5 min, based on the temporal profile of the effect. Time-course analyses were performed in successive bins spanning 5 min, 5-10 min, and 10-15 min.

#### Von Frey mechanical sensitivity testing

Mice were implanted with a chronic optical fiber (200 µm 0.39 NA; Thorlabs) targeting the vlPAG. Fibers were positioned using stereotaxic coordinates (relative to bregma) centered on the vlPAG at anterio-posterior (AP) = −4.5, medio-lateral (ML) = −1.2, and dorso-ventral (DV)= −3.2, with an angle of 20°.

Mechanical sensitivity was assessed using calibrated von Frey filaments via the up-and-down method (Chaplan et al., 1994). Mice were first habituated for 45 minutes to the experimental boxes and grid for two consecutive days before testing. Each mouse was then placed individually into a small compartment (6 × 12 cm) over a Von Frey mesh grid (Bioseb, France). Testing began with the 0.158 g filament applied perpendicularly to the plantar surface of the hind paw until the filament bowed slightly. Filaments of increasing or decreasing force (0.0045 g, 0.02 g, 0.068 g, 0.158 g, 0.178 g, 1.20 g, 2.041 g, and 5.495 g; Bioseb BIO-VF-M) were applied according to the subject’s response. A positive response was defined as clear, rapid paw withdrawal, licking, or flinching during application. Following the first response transition, testing continued for four additional trials. A positive response prompted the use of the next lower force filament, whereas a negative response prompted the next higher force filament. Left and right hind paws were tested independently at each time point (baseline, 30, 60 and 90 minutes post-injection). Data are presented as individual animal traces overlaid with the group mean ± SEM.

Systemic ABT594 (120 µg/kg) was administered i.p. and compared with saline (n = 6 per group). In the photopharmacology part, NV-ABT594 was administered i.p. at two doses (0.8 mg/kg and 4 mg/kg), with a saline control group. All injections were administered at 100 µl per 10 g body weight. Local photorelease of ABT594 was induced by delivering 365nm light pulses (1 second pulses at 0.1 Hz) through the implanted optical fiber in the vlPAG. Light stimulation was applied during the 30- and 60-minutes post-injection behavioral testing epochs. Mechanical thresholds were measured during illumination (365-nm light-on) as well as before illumination onset (365-nm light-off at baseline) and after illumination was discontinued (90 min) to determine whether behavioral effects were light-dependent and reversible. Saline-injected animals received identical 365-nm light stimulation to control for any non-specific effects of light delivery.

### GENERAL STATISTICAL ANALYSIS

Statistical analyses were performed using R (v4.6.0; RRID:SCR_001905) in RStudio (v2026.04.0+526) using custom scripts and the packages lme4, lmerTest, emmeans, drc, tidyverse and ggplot2, Python (v3.9; RRID:SCR_008394) via PyCharm CE (v2024.3.6.1; RRID:SCR_018221), GraphPad Prism 10 (RRID:SCR_002798) or MATLAB (RRID:SCR_001622). Statistical procedures, sample sizes, definitions of n, exact *p* values and multiple-comparison adjustments are reported in the corresponding figure legends. Unless otherwise indicated, tests were two-sided and the significance threshold was *p* < 0.05.

The biological experimental unit was an individual mouse. Repeated observations obtained from the same mouse, including photometry stimulation events, behavioral time bins, paw measurements and electrophysiological trials or units, were treated as nested or repeated measurements rather than as independent biological replicates. For repeated-measures datasets, we used linear mixed-effects models (LMMs) fitted with lme4, with mouse identity included as a random intercept at minimum. Fixed-effect significance was assessed using Type III analysis of variance with Satterthwaite approximation of degrees of freedom, implemented in lmerTest. Where justified by the experimental design, post-hoc contrasts were estimated using emmeans and adjusted for multiple comparisons using Dunnett, Tukey or Holm correction as specified for each analysis.

For datasets with a single independent observation per mouse and two groups, we used Welch’s t-tests. For data sets with two paired observations per mouse, we used Wilcoxon paired test, and if multiple groups were analyzed, we used effects analysis followed by Sidak’s multiple comparisons test. For one-sample comparisons, we used one-sample t-tests. When model assumptions were materially violated or sample sizes were insufficient to support reliable parametric inference, the corresponding non-parametric test was used as a sensitivity analysis, as specified in the figure legend.

Data are displayed as individual biological replicates whenever possible, with group summaries shown as mean ± SEM No statistical methods were used to predetermine sample sizes. Mice were assigned to experimental groups as described for each experiment, and data collection and analysis were randomized and/or blinded where feasible. Exclusion criteria were defined before analysis and are stated in the relevant Methods sections and figure legends.

## Acknowledgements

This work was supported by a Human Frontier Science Program Research Grant (RGP0035/2020 to AM and GED), by the Agence Nationale de la Recherche (ANR-24-CE16-6957 NIMONAR to AM and NR), the National Institutes of Health (NS111600 to GED and AM), the Major Research Program of PSL Research University ‘PSL-Neuro’ implemented by the ANR (ANR-10-IDEX-0001), the Fondation pour la Recherche Médicale (FRM EQU201903007961 and EQU202503019989 to P.F.) and the French National Cancer Institute (INCa grant SPAV1-23-005 to P.F.). This project has received funding from the European Union’s Horizon 2020 research and innovation programme under the Marie Skłodowska-Curie Grant Agreement n° 101023337 (to AB). JJ was supported by a Memolife Labex, fourth-year PhD fellowship. NG was supported by a Swedish Research Council (Vetenskapsrådet) International Post-doctoral fellowship (2022-06168).

We thank DIM C-BRAINS (Région Île-de-France) and NeuroPSL (Université PSL), Centre National de la Recherche Scientifique (CNRS) UMR 8249, as well as the ESPCI Paris for their support. We are grateful to our colleagues from the Neurophysiology and Behavior lab and the Brain Plasticity Unit for their valuable discussions and comments. We also thank Audrey Combès (Laboratory of Analytical, Bioanalytical Sciences and Miniaturization, Chemistry, Biology and Innovation (CBI), UMR 8231, ESPCI Paris) for her assistance with HPLC experiments, Aylin Gulmez and Noé Dubois-Thévenot for their help with immunohistochemistry experiments, Nicolas Debray for his work on the camera recording software, Yasmine Layadi for her help with setting up behavior experiments, Jean-Pierre Hardelin for his input on the manuscript, and Deniz Dalkara (Institut de la Vision, Paris) for viral production. Additionally, we acknowledge Hélène Geoffroy and Jeldy Cubas-Hernandez for their administrative support, Steve Didienne and the ESPCI Paris workshop for their technical assistance in constructing the RTPP setups, as well as Otilia de Oliveira, Lizandra Arencibia Sorzano, and Emilie Tubeuf for their care of the animal facility. Images of mice in Fig. 1a, 2d, 3a, 4a, 5a,d and 6c were adapted from SciDraw (https://scidraw.io/).

## Author contributions

NG, GED and AM designed research

NG, MM, AB, GG, JJ, NP, AF, NR and GED performed research

NG, MM, AB, GG and NR analyzed data

AF, PF, GED and AM supervised research

NG and AM wrote the paper, with inputs from all the authors

## Supplementary material

**Figure S1:**
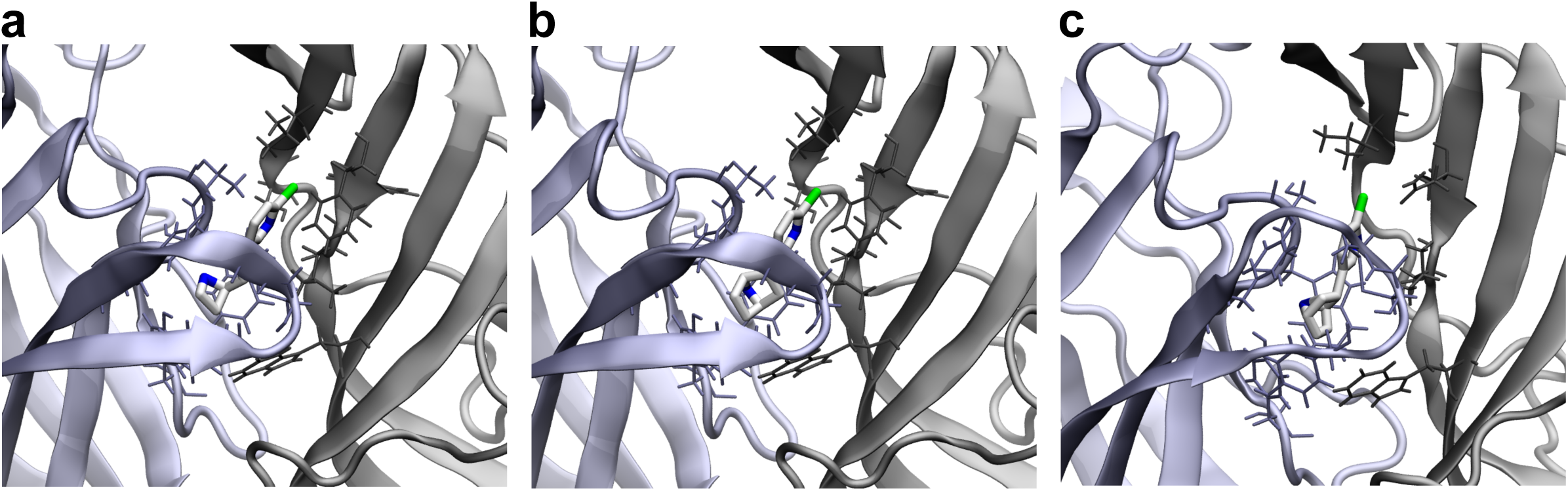
Structural modeling of NV-caged nicotinic agonists. **a**, Predicted binding pose of ABT594 in the orthosteric binding site of the α4β2 nAChR (PDB 8ST0). CDocker interaction energy: −38.9 kJ/mol. Receptor residues interacting with ABT594 are shown. NV-ABT594, docked under identical conditions, failed to produce productive binding orientations, generating only aberrant poses with unfavorable scores that are not meaningful in the absence of rational ligand placement. **b**, Predicted binding pose of epibatidine in the orthosteric binding site of the α4β2 nAChR (PDB 8ST0). CDocker interaction energy: −41.0 kJ/mol. Interacting receptor residues are shown. NV-EPI, docked under identical conditions, failed to produce productive binding orientations, generating only aberrant poses with unfavorable scores that are not meaningful in the absence of rational ligand placement. **c**, High-resolution structure of the α7 nAChR in complex with epibatidine (PDB ID 7KOQ).

**Figure S2:**
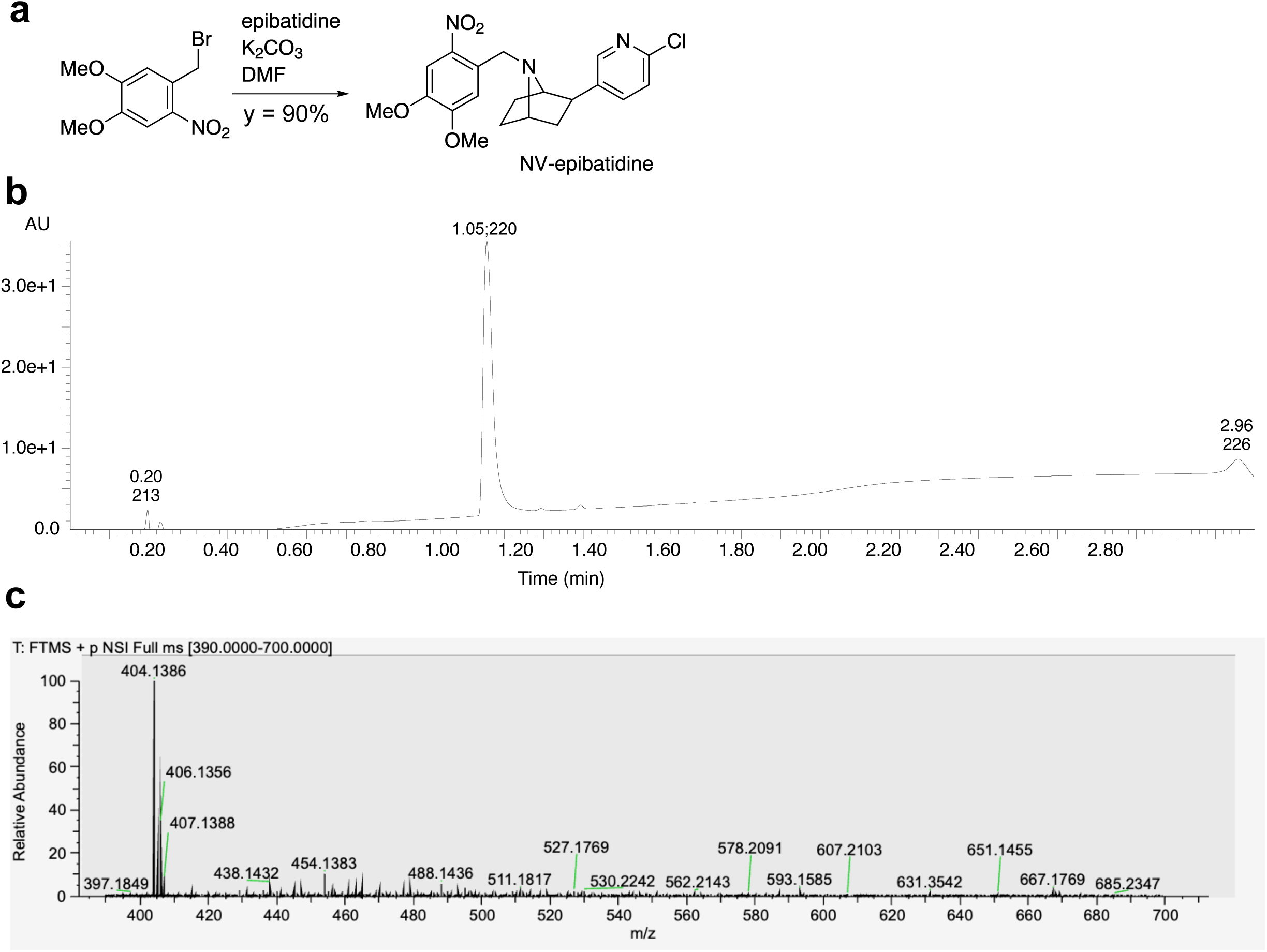
Synthesis and chemical characterization of NV-EPI. **a**, Synthesis scheme of NV-EPI. **b**, UPLC chromatogram of purified NV-EPI. **c**, ESI mass spectrogram of NV-EPI.

**Figure S3:**
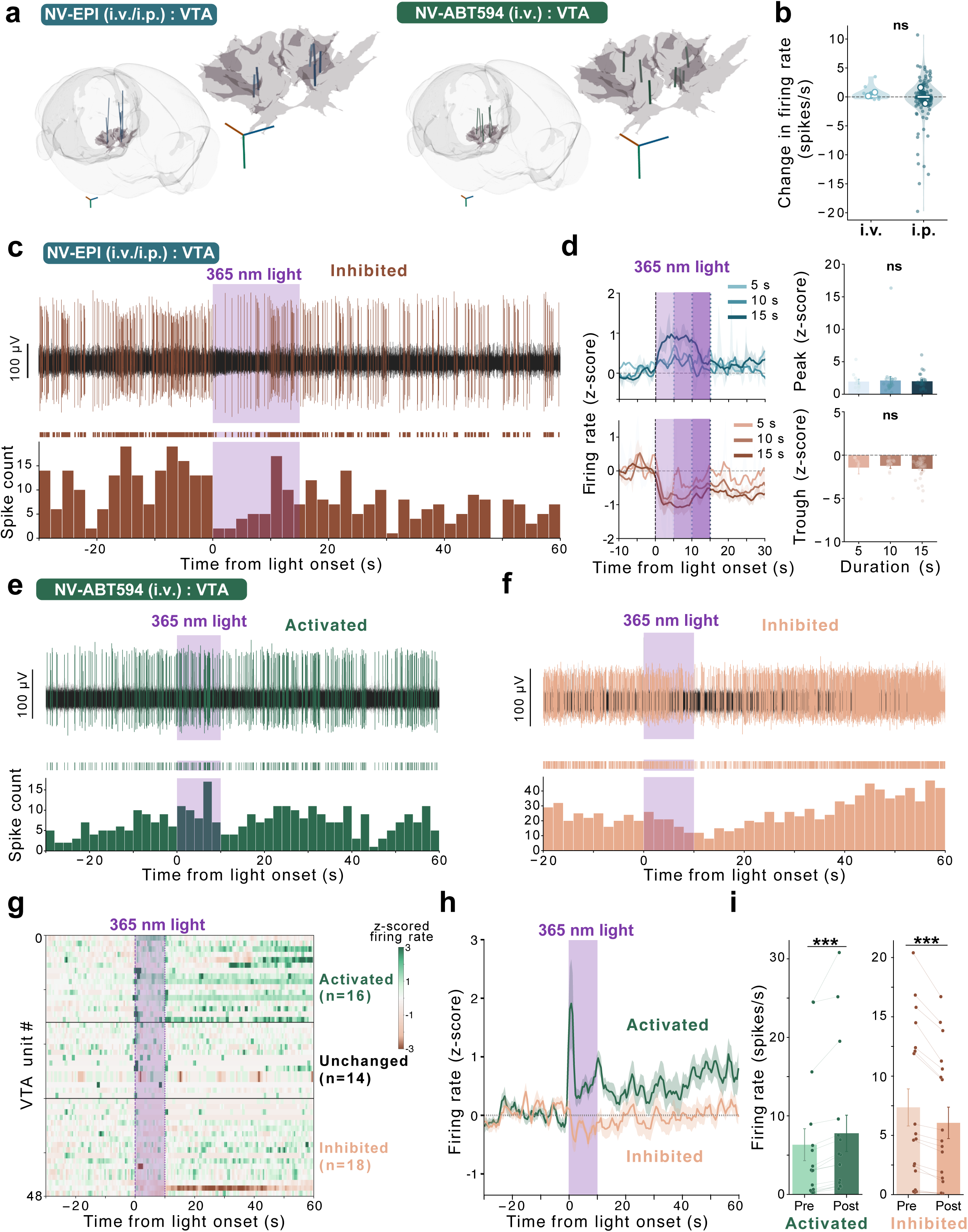
Light-evoked responses in the VTA **a,** Three-dimensional reconstruction of silicon-probe trajectories targeting the ventral tegmental area. The ventral tegmental area (VTA) is highlighted as a translucent volume. Close-up views show the VTA and the distal 315 µm recording span of each probe, corresponding to the 32 recording channels. Each trajectory color represents an individual silicon-probe track. The three-dimensional scale bars indicate 500 µm along the anteroposterior (orange), mediolateral (blue), and dorsoventral (green) axes. **b**, Light-evoked firing-rate changes following 15-s illumination are shown for VTA units after intravenous (i.v.) or intraperitoneal (i.p.) NV-EPI administration. Filled points represent individual units, open circles indicate mouse means, violins show the unit-level distributions, thick vertical lines denote the interquartile range and white horizontal lines indicate the median. Administration route was not significantly associated with response amplitude (i.v., n = 11 units from two mice; i.p., n = 132 units from two mice; adjusted i.p. – i.v. difference, −0.43 Hz; 95% CI, −4.02 to 3.17; *p* = 0.8). Effects were estimated using linear mixed-effects models with mouse as a random intercept and adjustment for time from injection to illumination. These results supported pooling i.v.- and i.p.-administered NV-EPI recordings in the subsequent analyses of light-evoked responses. **c**, Representative extracellular voltage trace (top) and spike-rate histogram (bottom, 2-s bins) from a VTA unit inhibited by a 365-nm light stimulus (15 s, 8 mW) after systemic NV-EPI administration. Shading indicates light delivery. Related to Fig. 2g-i. **d**, Effect of illumination duration on VTA responses to photorelease of EPI. Left, mean ± SEM z-scored firing rate for activated and inhibited units across 5-, 10-, and 15-s illumination durations, aligned to light onset. Right, peak responses of activated units and trough responses of inhibited units. Response amplitude did not differ significantly among illumination durations for activated units (n = 41 units from five mice; likelihood-ratio test, χ²(2) = 2.60, *p* = 0.3;) or inhibited units (n = 44 units from four mice; χ²(2) = 2.07, *p* = 0.3). **e**, Representative extracellular voltage trace (top) and spike-rate histogram (bottom) from a VTA unit activated by 365-nm light after systemic NV-ABT594 administration. **f**, As in **e**, for a light-inhibited VTA unit. **g**, Heatmap of trial-averaged, z-scored firing rates for VTA units recorded after systemic NV-ABT594 administration (4 mg/kg), aligned to light onset (10 s, 8 mW). Each row represents one unit averaged across three trials. Units are ordered by response classification as activated (n = 16), unchanged (n = 14) or inhibited (n = 18); n = 48 units from six mice. **h**, Mean ± SEM z-scored firing rate over time for activated (n = 16) and inhibited (n = 18) VTA populations following NV-ABT594 uncaging, aligned to light onset. **i**, Firing rates before and after light onset for individual activated (left; n = 16) and inhibited (right; n = 18) VTA units following NV-ABT594 uncaging. Paired points represent individual units; bars show mean ± SEM. Firing rate increased in activated units by 1.45 spikes/s (95% CI, 0.72 to 2.19; linear mixed-effects model, FDR-adjusted *p* < 0.001) and decreased in inhibited units by 1.31 spikes/s (95% CI, −1.80 to −0.82; FDR-adjusted *p* < 0.0001).

**Figure S4:**
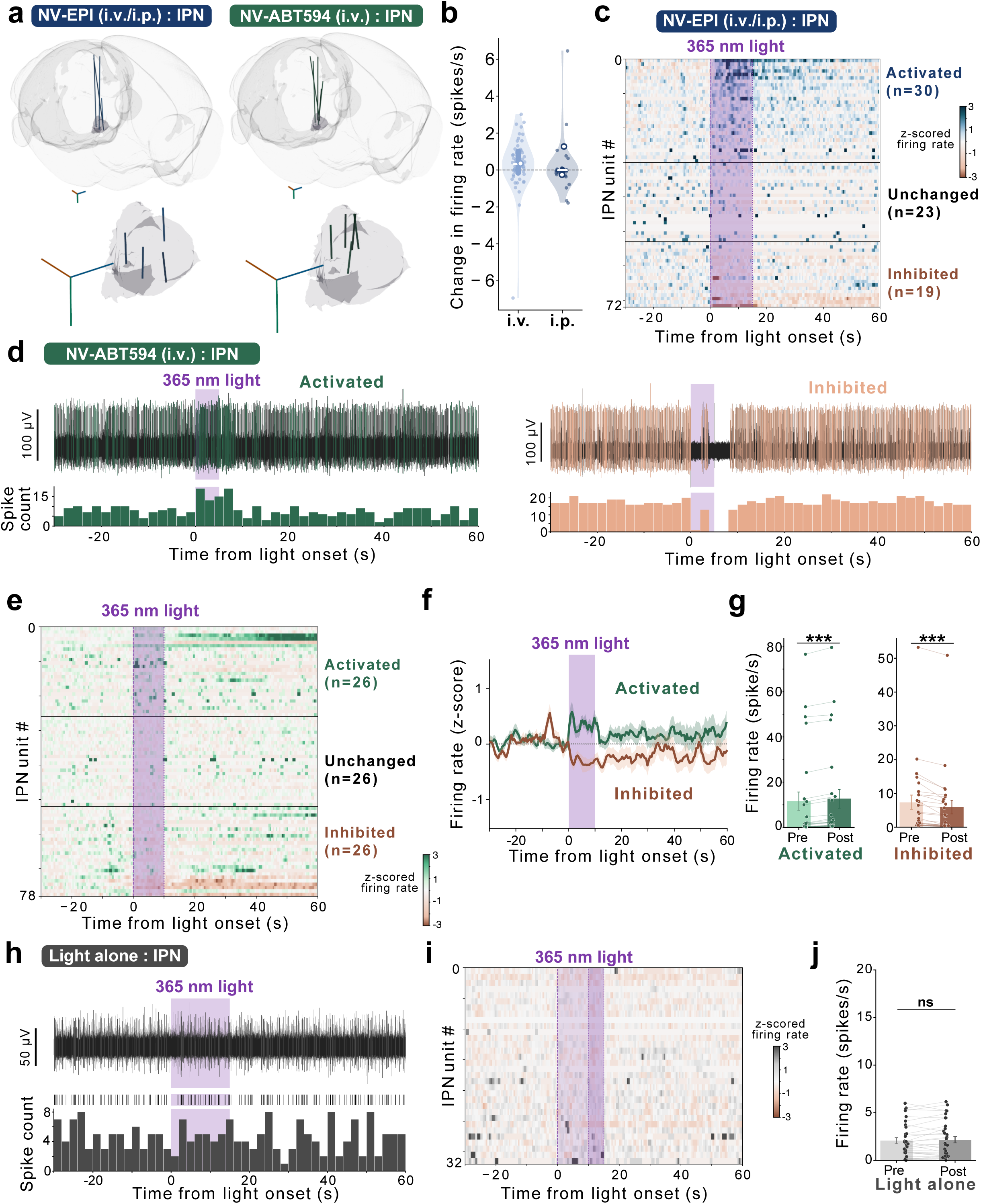
Light-evoked responses in the IPN, and light-alone controls **a**, Three-dimensional reconstruction of silicon-probe trajectories targeting the interpeduncular nucleus. The interpeduncular nucleus (IPN) is highlighted as a translucent volume. Close-up views show the IPN and the distal 315 µm recording span of each probe, corresponding to the 32 recording channels. Each trajectory color represents an individual silicon-probe track. The three-dimensional scale bars indicate 500 µm along the anteroposterior (orange), mediolateral (blue), and dorsoventral (green) axes. **b**, Light-evoked firing-rate changes following 15-s illumination are shown for IPN units after intravenous (i.v.) or intraperitoneal (i.p.) NV-EPI administration. Filled points represent individual units, open circles indicate mouse means, violins show the unit-level distributions, thick vertical lines denote the interquartile range and white horizontal lines indicate the median. Administration route was not significantly associated with response amplitude (i.v., n = 51 units from one mouse; i.p., n = 34 units from two mice; adjusted i.p. – i.v. difference, −1.05 Hz; 95% CI, −8.08 to 5.97; p = 0.8). Effects were estimated using linear mixed-effects models with mouse as a random intercept and adjustment for time from injection to illumination. These results supported pooling i.v.- and i.p.-administered NV-EPI recordings in the subsequent analyses of light-evoked responses. **c**, Heatmap of mean z-scored firing rate for all recorded IPN units following systemic NV-EPI (n = 72 units from 3 mice), aligned to the onset of the 15-s light stimulus and averaged across three trials, sorted into activated (n = 30), unchanged (n = 23), and inhibited (n = 19) populations. **d**, Representative voltage traces and spike-rate histograms (spikes/2 s bins) from an activated (left) and an inhibited (right) IPN neuron following systemic NV-ABT594, aligned to 365-nm light onset. **e**, Heatmap of mean z-scored firing rate for all recorded IPN units following systemic NV-ABT594 (n = 78 units from 4 mice), aligned to 365-nm 10-s light onset and averaged across three trials, sorted into activated (n = 26), unchanged (n = 26), and inhibited (n = 26) populations. **f**, Mean ± SEM z-scored firing rate over time for activated and inhibited IPN populations following NV-ABT594 uncaging, aligned to light onset. **g**, Firing rates before and after light onset for individual activated (left) and inhibited (right) IPN units following NV-ABT594 uncaging; bars show mean ± SEM. For activated units, firing rate increased during light delivery by 1.083 spikes/s (95% CI, 0.763 to 1.403, LMM FDR-adjusted *p* < 0.0001), whereas inhibited units decreased their firing by 1.382 spikes/s (95% CI, −1.908 to −0.856; FDR-adjusted *p* < 0.0001). **h**, Representative extracellular voltage trace and spike-rate histogram (spikes/2 s bins) from an IPN neuron during light delivery alone (no NV-EPI), showing no detectable light-locked change in firing. **i**, Heatmap of trial averaged, z-scored firing rate for recorded IPN units aligned to the onset of 10- or 15-s light stimuli. and averaged across three trials. None of the 32 units met the prespecified criteria for activation or inhibition (n = 32 units from 3 mice) **j**, Firing rates before and during light delivery for individual IPN units in light alone control experiments. Individual paired points show IPN light alone unit observations from 10-s and 15-s illumination protocols; bars indicate mean ± SEM. The mean change in firing rate was 0.050 spikes/s (90% CI, −0.179 to 0.278; n = 32 units from 3 mice). The 90% CI was contained within the prespecified ±0.5 spikes/s equivalence bounds, supporting equivalence to a negligible average effect (two one-sided tests *p* < 0.001).

**Figure S5:**
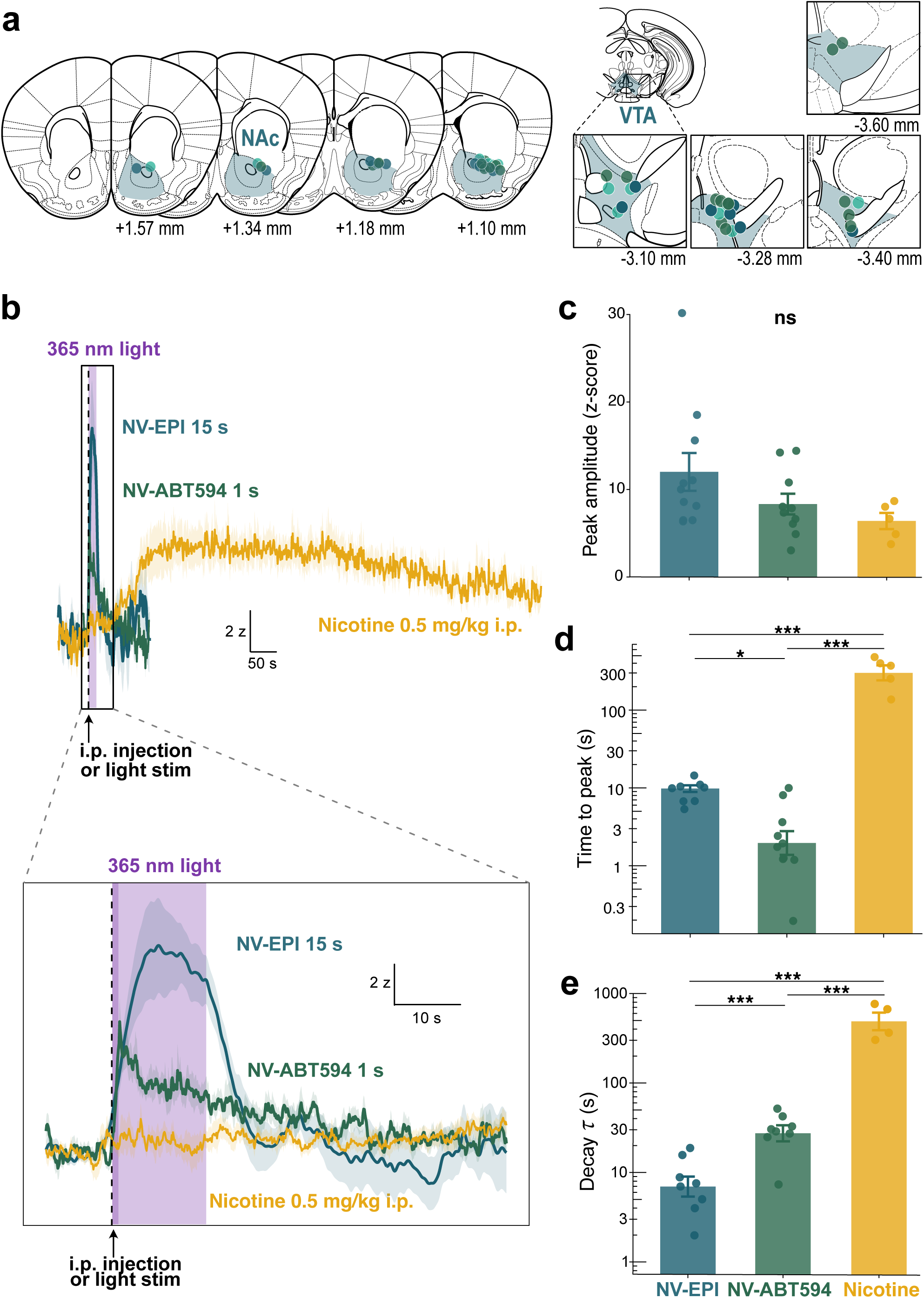
Comparison of NAc dopamine dynamics evoked by local photorelease of nicotinic agonists and systemic nicotine **a,** Anatomical reconstructions of optical-fiber placements in the NAc and VTA. Optical-fiber tracks are shown for mice used with NV-Epi only in teal, mice used with NV-ABT594 only in green, and mice used with both compounds in turquoise. Reconstructions were available for 25 of 34 NAc fiber tracks and for 24 of 34 VTA fiber tracks. AP coordinates are given in millimeters relative to bregma. **b**, Left, average NAc DA responses following systemic nicotine (i.p. 0.5 mg/kg, n = 5 mice), ABT594 uncaging (1-s light, injected i.p. at 4 mg/kg, n = 10 mice) or EPI uncaging (15-s light, injected i.p. at 200 μg/kg, n = 11 mice), aligned to injection or light stimulation onset. Right, zoom on the 60 first seconds of the response (inset on the left). Solid lines, group means; shaded areas, SEM **c**, Peak amplitude (z-score) for the same three conditions. NV-EPI: 12.0 ± 2.2; NV-ABT594: 8.3 ± 1.2; Nicotine: 6.4 ± 0.9. Peak amplitudes did not differ significantly across conditions (one-way ANOVA, F(2,23) = 2.30, p = 0.12; all Tukey *p* > 0.15). Bars show group means ± SEM; individual points show single mice. **d**, Time to peak (s, log10 scale) for the same three conditions. NV-ABT594 peaked fastest (3.2 ± 1.0 s), followed by NV-EPI (10.3 ± 0.9 s; Tukey *p* = 0.04), while nicotine peaked ∼30 times slower (328.4 ± 60.3 s; LMM, F(2,2.0) = 78.4, *p* = 0.01; EPI vs Nic p < 0.0001, ABT vs Nic *p* < 0.0001). Bars show geometric means, with error bars representing ± 1 SEM in log space; individual points represent mice. The y-axis is logarithmic (log10). **e**, Decay time constant (τ, s) for the same three conditions. NV-EPI: 8.6 ± 2.1 s (n = 8); NV-ABT594: 30.8 ± 4.6 s (n = 8); nicotine: 525 ± 113 s (n = 4). Decay τ differed across conditions (LMM, *F*(2,17) = 159,407, *p* < 0.0001; NV-EPI versus nicotine, Tukey *p* < 0.0001; NV-ABT594 versus nicotine, Tukey *p* < 0.0001). The ∼60-fold longer decay τ for nicotine reflects its prolonged temporal profile relative to the rapid photopharmacological DA transients. Bars show geometric means, with error bars representing ± 1 SEM in log space; individual points represent mice. The y-axis is logarithmic (log10).

**Figure S6.**
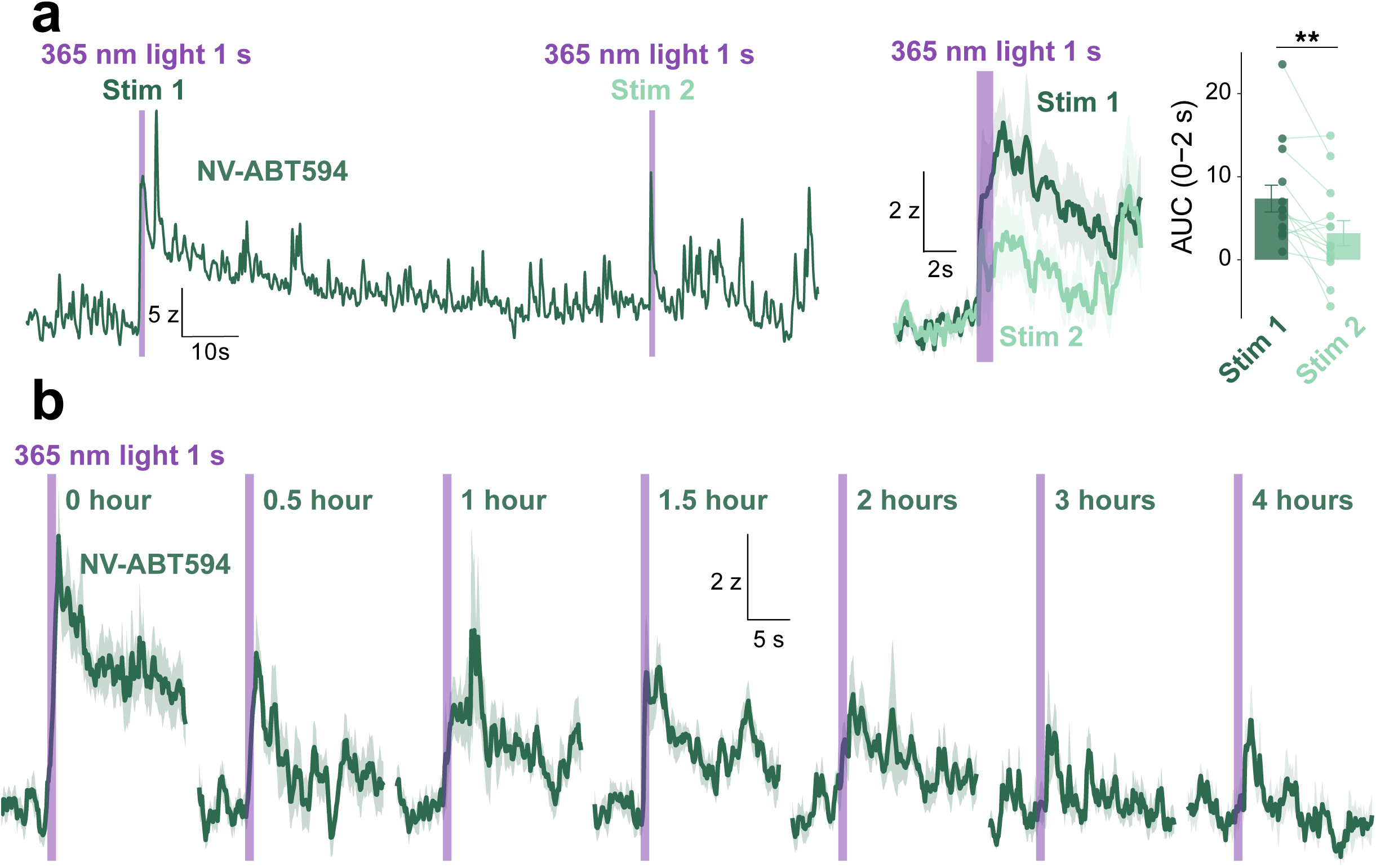
Repeatability and temporal availability of NV-EPI and NV-ABT594 uncaging. **a**, Left, representative trace showing responses to two consecutive 1-s violet-light stimulations; violet shading indicates light delivery. Right, traces (mean ± SEM) and paired AUC values for the first and second stimulations. AUC was lower for Stim 2 than Stim 1 (3.20 ± 1.51 versus 7.36 ± 1.63 z·s, mean ± SEM; linear mixed-effects model with mouse as a random effect, *p* = 0.004). Bars show mean ± SEM; 14 recordings from 10 mice. **b**, Repeatability of NV-ABT594 uncaging over hours. Averaged traces (mean ± SEM) aligned to 1s violet stimulations delivered 0, 0.5, 1, 1.5, 2, 3, and 4h after NV-ABT594 injection (4 mg/kg, i.p.; n = 10 mice).

**Figure S7:**
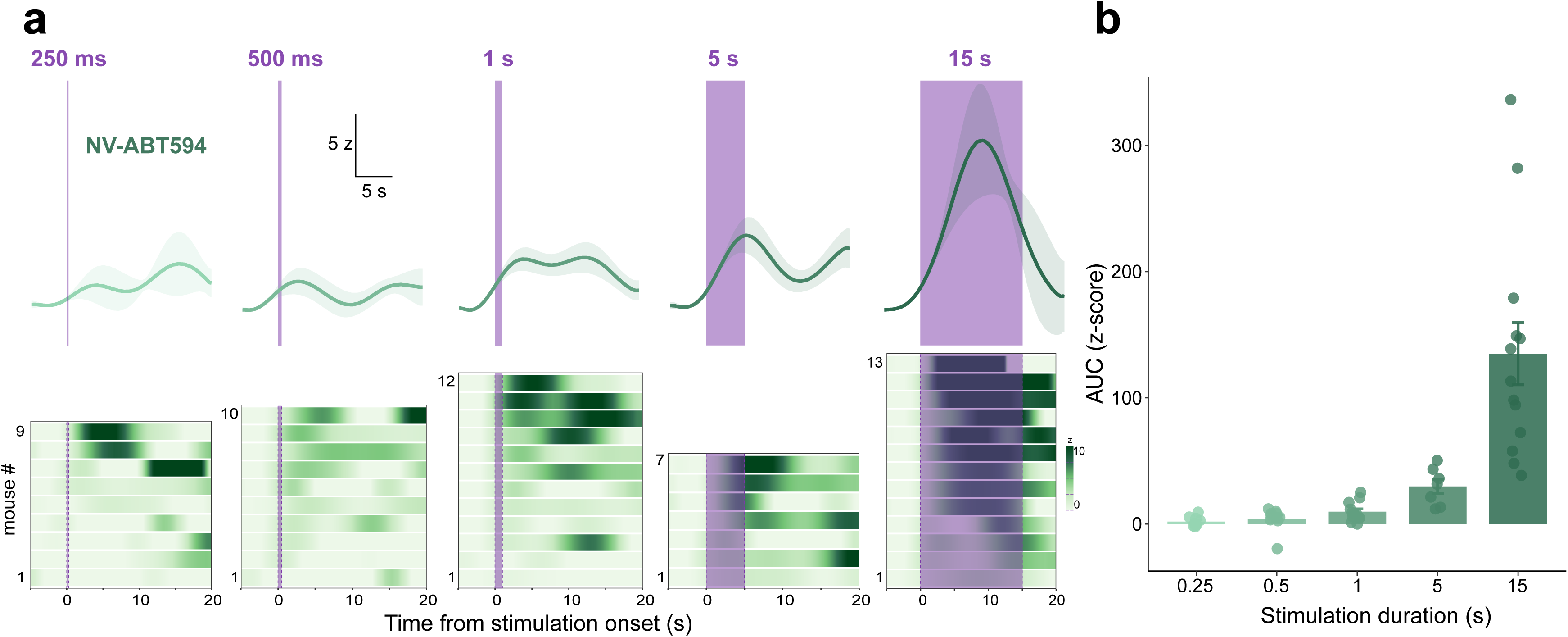
Illumination-duration dependence of NV-ABT594-evoked DA responses in the NAc. **a,** Mean (± SEM) traces (top) and per-mouse heatmaps (bottom) showing NAc DA responses to VTA NV-ABT594 uncaging at increasing illumination durations (250 ms, 500 ms, 1 s, 5 s, 15 s; violet shading indicates light delivery), aligned to stimulation onset. n = 9, 10, 12, 7, and 13 mice, respectively. **b,** Quantification of response magnitude as AUC of the z-scored DA response, integrated from stimulation onset until 3 s after stimulation offset (corresponding to windows of 0-3.25, 0-3.5, 0-4, 0-8 and 0-18 s, respectively). Bars show group mean ± SEM; individual points show per-mouse means (n = 7-13 mice per duration, 17 mice total). AUC increased with illumination duration (linear mixed-effects model with mouse as a random intercept: *F*(4,13.38) = 19.32, *p* < 0.0001). Modelling duration as a continuous predictor confirmed a strong positive relationship between illumination duration and AUC (*β* = 8.28 ± 0.90 z-score per second of illumination, *t*(31.78) = 9.19, *p* < 0.0001). Dunnett-adjusted comparisons against the 250 ms condition showed a significantly larger response at 15 s (*p* < 0.0001).

**Figure S8.**
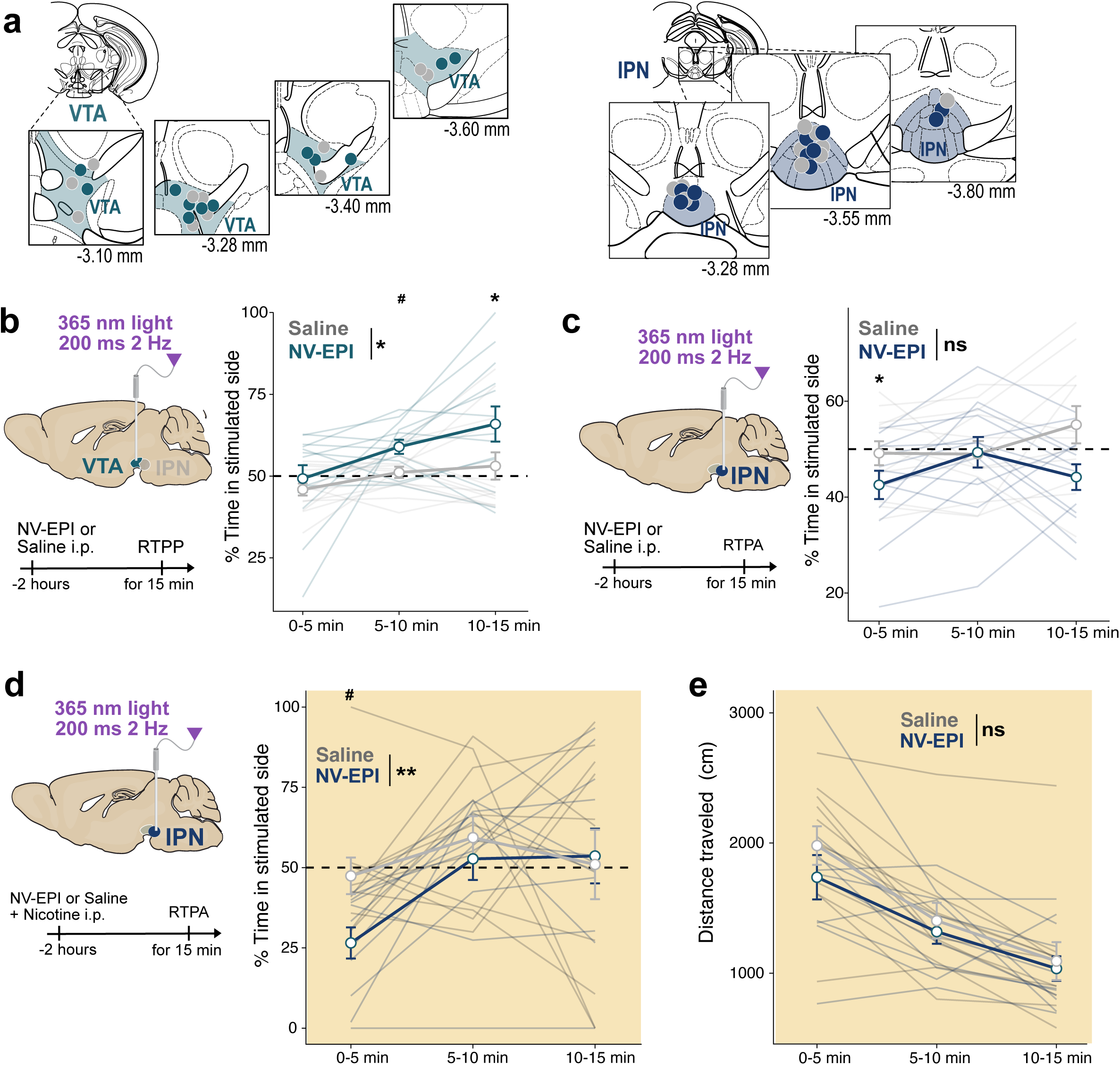
Time-resolved dynamics of place preference and aversion. **a,** Anatomical reconstructions of optical-fiber placements in the VTA and IPN. Placements are shown for NV-EPI mice in color and saline mice in gray. Reconstructions were available for 23 of 26 VTA-implanted mice and all 25 IPN-implanted mice. **b**, Time-resolved percent time in the stimulated side across three 5-minute bins (0-5, 5-10, 10-15 min) for the VTA RTPP experiment (saline vs. NV-EPI), corresponding to Fig. 5e. Mixed-effects ANOVA revealed significant main effects of Group (*F*(1,24) = 6.24, *p* = 0.02, ηG²=0.098) and TimeBin (*F*(1.39,33.42) = 6.73, *p* = 0.008, ηG²=0.141), with no significant Group×TimeBin interaction (*F*(1.39,33.42) = 1.05, *p* = 0.3). NV-EPI mice differed from saline controls during the 5–10-minute bin (*p* = 0.03) and spent significantly more time in the stimulated side than chance during this period (t(12) = 4.14, *p* = 0.001, Bonferroni-adjusted *p* = 0.008). **c**, Time-resolved percent time in the stimulated side across three 5-minute bins for the naive-mouse IPN RTPA experiment (saline vs. NV-EPI), corresponding to Fig. 5g. A two-way repeated-measures ANOVA with group as the between-subject factor revealed a significant group-by-time-bin interaction (*F*(2,46) = 3.83, *p* = 0.03, generalized η² = 0.049), with no significant main effects of group (*F*(1,23) = 2.59, *p* = 0.1) or time bin (*F*(2,46) = 1.99, *p* = 0.15). Within the NV-EPI group, occupancy was significantly lower during the first 5 minutes than during the 5–10-minute bin (Bonferroni-adjusted *p* = 0.02). One-sample tests against 50% chance occupancy showed that NV-EPI mice were not significantly below chance during any time bin after correction. **d**, Left, time-resolved percent time in the stimulated side for the nicotine-primed IPN RTPA experiment (saline vs. NV-EPI), corresponding to Fig. 5h. During the first 5 min, NV-EPI mice spent 26.5 ± 4.9% of time in the stimulated side, significantly below 50% chance occupancy (Wilcoxon signed-rank test, *W* = 0, *p* = 0.004, Bonferroni-adjusted *p* = 0.02), whereas saline-treated mice did not differ from chance (47.4 ± 5.7%; *W* = 14, adjusted *p* = 0.6). NV-EPI mice also spent significantly less time in the stimulated side than saline-treated mice during the first 5 minutes (Bonferroni-adjusted *p* = 0.03), with no significant between-group differences during subsequent bins. **e**, Locomotor distance traveled for the nicotine-primed IPN RTPA experiment (saline vs. NV-EPI), corresponding to Fig. 5h, did not differ between groups (Group: *F*(1,20) = 0.657, *p* = 0.4; Group×TimeBin: *F*(1.41,28.26) = 0.605, *p* = 0.5; between-group comparisons at each bin, all *p* ≥ 0.29).

---

**Supplementary Movie 1.** fUSi during photorelease. **Top,** Each voxel represents means cerebral blood volume (CBV) after realignment and averaging over the first six trials during photorelease (left) and over the first two trials during control stimulation (right). Time is relative to light stimulation onset. Note the strong long-lasting unilateral vascular activation following photo release in a region encompassing the stimulated VTA. **Bottom**, CBV time series after realignment and averaging over the first 6 trials during photorelease (left) and during control stimulation (right) in the stimulated vs contralateral VTA (left), and in the stimulated VTA during photo release vs control (right). Violet shaded box: light stimulation.

